# Differential regulation of local mRNA dynamics and translation following long-term potentiation and depression

**DOI:** 10.1101/2020.07.08.192369

**Authors:** Paul G Donlin-Asp, Claudio Polisseni, Robin Klimek, Alexander Heckel, Erin M Schuman

## Abstract

Decades of work have demonstrated that mRNAs are localized and translated within neuronal dendrites and axons to provide proteins for remodeling and maintaining growth cones or synapses. It remains unknown, however, whether specific forms of plasticity differentially regulate the dynamics and translation of individual mRNA species. To address these issues, we targeted three individual synaptically-localized mRNAs, *CamkIIa*, *Beta actin*, *Psd95*, and used molecular beacons to track endogenous mRNA movements and reporters and Crispr-Cas9 gene editing to track their translation. We found widespread alterations in mRNA behavior during two forms of synaptic plasticity, long-term potentiation (LTP) and depression (LTD). Changes in mRNA dynamics following plasticity resulted in an enrichment of mRNA in the vicinity of dendritic spines. Both the reporters and tagging of endogenous proteins revealed the transcript-specific stimulation of protein synthesis following LTP or LTD. The plasticity-induced enrichment of mRNA near synapses could be uncoupled from its translational status. The enrichment of mRNA in the proximity of spines allows for localized signaling pathways to decode plasticity milieus and stimulate a specific translational profile, resulting in a customized remodeling of the synaptic proteome.

## Introduction

Synaptic plasticity requires the rapid and robust remodeling of the proteome_1_. Both the strengthening (long term potentiation-LTP) and the weakening of synaptic connections (long term depression-LTD) requires proteome remodeling_2_. Neurons use diverse mechanisms to achieve this regulation including the posttranslational modifications of proteins_2_, transcriptional changes_3_, and translational changes_4_. Indeed, neurons can rapidly regulate and control synaptic proteomes by localizing and translating of mRNAs in axons and dendrites_5–12_. A number of key synaptic proteins are encoded by translationally regulated mRNAs including ARC_13–15_, FMRP_16_, PSD-95_16,17_, and CAMK2a_18_. Given the capacity for protein synthesis in distal compartments, a fundamental question is how dendritically and axonally localized mRNAs become recruited near synapses and then translationally regulated locally during plasticity.

Current evidence suggests that single mRNAs, bound by RNA-binding proteins (RBPs), interact with the cytoskeleton for long distance transport from the cell body to the dendrites and axons_19,20_. Both localization_19_ and translational regulatory elements_21_ are present in the 5’ and 3’ untranslated regions (UTRs) of mRNAs. Trans-acting factors, including RBPs and microRNAs (miRNAs), interact with UTR elements to regulate mRNA localization and translation; these interactions are also regulated by plasticity. mRNAs are believed to be transported in a translationally quiescent state-likely only engaged in translation near synapses_22,23_. The dynamic and bidirectional_22,24–26_ scanning behavior of mRNAs in dendrites allows, in principle, for the capture and translation of mRNAs as needed for proteome maintenance and remodeling_22,23_. Our understanding of these processes, however, is largely derived from live imaging experiments for a limited number of individual candidate mRNAs *Beta actin*_22,25_ and *Arc*_27_; the relationship between the sequestration/capture of RNAs and their translation during plasticity is not well understood.

To address this, we used molecular beacons to track and quantify the dynamics of 3 endogenous mRNAs under basal conditions and after plasticity. We found that induction of either long-term potentiation (LTP) or metabotropic glutamate receptor long-term depression (mGluR-LTD) resulted in widespread attenuation of mRNA motility and led to an enrichment of mRNA near dendritic spines. These enhanced mRNA dynamics and availability near synapses was accompanied for some, but not all, mRNAs by enhanced translation of either a reporter or a CRISPR/Cas9 tagged endogenous protein. This dissociation allows for the enrichment of mRNAs near spines where localized signaling pathways can control which specific sets of transcripts are translated.

## Results

### Tracking endogenous mRNA dynamics in live neurons

To assess endogenous mRNA dynamics, we focused on *Camk2a*, *Beta actin* and *Psd95* as they are abundant in neuronal dendrites_28_ and are translationally regulated by plasticity_18,22,29,30_. To track these mRNAs, we employed molecular beacons (Figure 1A)_31_. These mRNA-specific complementary oligonucleotides bear both a fluorophore and a quencher; the binding of a beacon to its targeted mRNA separates the fluorophore and quencher, resulting in a fluorescent signal that can be tracked in live cells. Similar probes have recently been used to track endogenous *Beta actin* in Xenopus axons_32_, yielding dynamic properties similar to those observed *in vivo* using the MS2-*Beta actin* mouse_25_.

**Figure 1:**
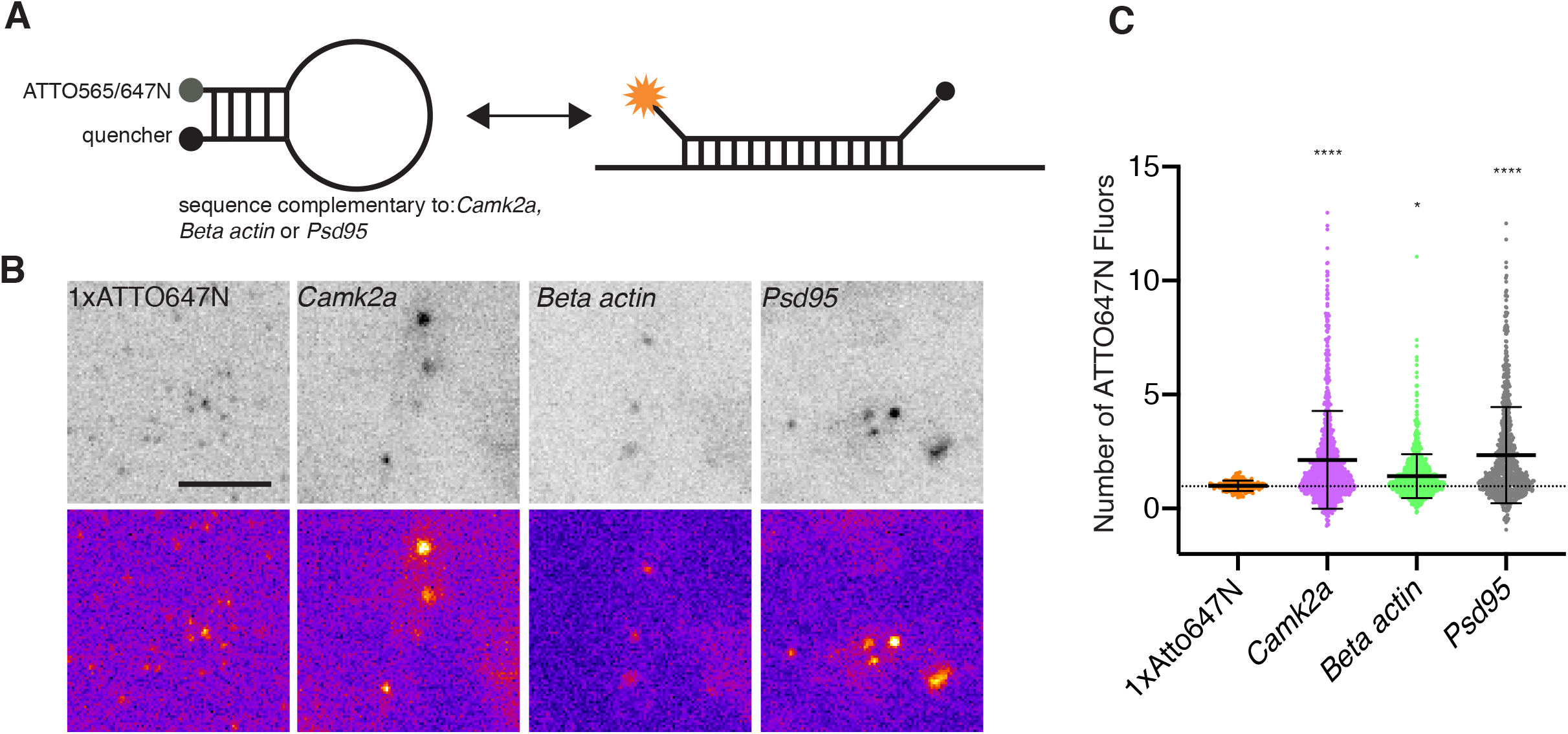
Dendritic mRNAs exist within transcript specific heterogeneous states within live neuron. **A.** Scheme of the molecular beacon design used in this study. A 5 nucleotide (nt) stem, with a reporter fluorophore on one side and a quencher dye on the other side were linked with a 26nt complementary sequence to *Camk2a*, *Beta actin* or *Psd95* mRNAs. When not hybridized to a target mRNA, the stem holds the reporter and quencher in proximity, preventing fluorescence. Upon binding to a target mRNA, the stem loop opens, and fluorescence can be detected in a reversible manner. **B.** Example images of Gatta quant standard and beacon labeled neurons shows a heterogenous distribution of particle intensities in live neurons. Scale bar = 5um. Data shown acquired at 5% laser power. **C.** Quantification of mRNA organizational state within dendrites shows a transcript-specific heterogenous distribution pattern of copy numbers of mRNAs per granule/puncta. 1x Atto647N (3/150); *Camk2a* (19/921), *Beta actin* (21/1003), *Psd95* (18/829) (cells/number of individual mRNA granules analyzed). Comparison relative to 1xATTO647N, Dunnett’s multiple comparison test: */**** p<.05/.0001. Data shown acquired at 5% laser power.

In primary rat hippocampal neuronal cultures (DIV21+) molecular beacons targeting endogenous *Camk2α*, *Beta actin* or *Psd95* mRNA (see methods) were imaged to report on endogenous mRNA dynamics for up to 20 min (Supplemental Videos 1-3). Interestingly, we observed a heterogenous size distribution for the mRNA puncta - with larger pronounced particles seen near the soma (Supplemental videos 1-3), similar to what has been reported previously for *Beta actin*_25_. In addition, we detected a number of apparent dendritic mRNA-mRNA fusion events (Supplemental Figure 1A, Supplemental Video 4), suggesting that these mRNAs can exist in a heterogenous copy number state-contrary to proposed models of single mRNA transport in axons and dendrites_29,32,33_. To assess this quantitatively, we analyzed the intensity of the individual beacon puncta (i.e. mRNA granules) and compared it to a commercially synthesized oligonucleotide containing a single ATTO647N fluorophore (see methods). Using this standard, we determined that a sizeable fraction of each mRNA exhibited an intensity consistent with a single mRNA species (Figure 1B&C, Supplemental Figure 1B&C). *Beta actin* mRNAs, in particular, were often detected in a range consistent with a single copy number state (Figure 1C, Supplemental Figure 1B&C), in line with previous reports_29,32_. Interestingly, while both *Camk2a* and *Psd95* mRNAs also existed as a single copies, a large fraction of the population existed in a multimeric state (Figure 1C, Supplemental Figure 1B&C), indicating that higher-order (containing more than a single mRNA) transporting mRNA granules exist within the dendrite. Furthermore, the relative abundance of either a single or multimeric state appears to be a transcript-specific feature-likely determined through specific sets of RNA-binding proteins bound to particular mRNAs.

To capture the dynamic behavior of each mRNA species within dendrites, we employed a semi-automated tracking approach. Using a custom-written analysis pipeline (see methods) we quantified the beacon mRNA dynamics (Figure 2A-C), distance traveled (Figure 2D) and transport velocity (Figure 2E) for all 3 mRNA targets. To assess mRNA dynamics, we measured the percentage of time (during the entire imaging epoch) a detected granule spent actively moving within the dendrite (either anterogradely: away from the cell body, or retrogradely: towards the cell body) or exhibited a confined behavior. For all experiments we imaged at 1 frame per second for up to 20 minutes (see methods). Confined behavior was assigned to periods of time when an mRNA granule exhibited restricted (< 0.5um) movement within the dendrite (see methods). Similar to previous reports_22,32_, we found that all 3 mRNAs spent most of their time in a confined state (*Camk2a*: 0.59+/− 0.10; *Beta actin*: 0.56 +/− 0.11; *Psd95*: 0.50 +/− 0.14. Mean fraction of time, +/− SEM). For active movement all three mRNAs displayed a slight bias for anterograde transport (anterograde vs. retrograde: *Camk2a*: 0.23 +/− 0.06 vs 0.18 +/− 0.07; *Beta actin*: 0.23 +/− 0.09 vs 0.22 +/− .06; *Psd95*: 0.31 +/− 0.11 vs 0.20 +/− 0.09) (Figure 2A), explaining how mRNAs can eventually populate more distal regions of the dendrite. Interestingly, *Psd95* granules exhibited a significant increased motility compared to *Beta actin* and *Camk2α.* All three mRNAs traveled similar distances (~20um on average) over the imaging epoch and exhibited similar velocities (~1um/s) for both anterograde and retrograde transport, consistent with the mixed polarity of microtubules within the dendrite_34_. We noted that while the majority of mRNA molecules we measured (orange and magenta highlighted puncta, Figure 2A&B) alternated between periods of confined vs. active transport, a small fraction of particles (green) showed little to no active transport during the entire imaging session. We therefore further distinguished confined vs truly stationary events (Supplemental Figure 2A, see methods) and found ~6% of the confined population were better characterized as stationary events. For all downstream analyses, we removed these stationary events from the analysis.

**Figure 2:**
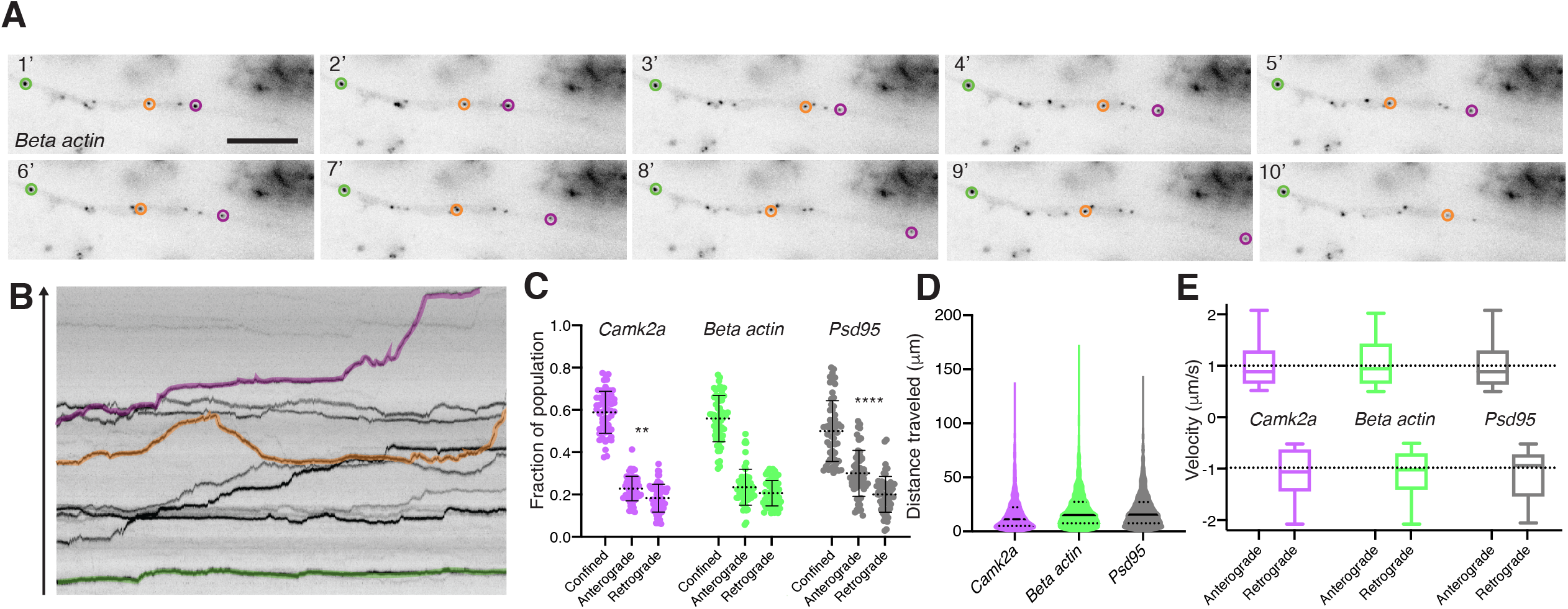
Tracking and classifying mRNA dynamics along dendrites in live neurons. **A.** Still images from a *Beta actin* 647N labeled dendrite (Supplemental video 5), shown are single frames every minute for 10 minutes (out of 20 total). Individual mRNA puncta are highlighted to illustrate distinct dynamic profiles, mainly stationary (green) or confined with periods of motility (orange and magenta). Scale bar = 5um. **B.** Kymograph from the first 10 minutes of (A/Supplemental video 5). Arrow denotes the anterograde direction along the dendrite. **C.** Quantification of mRNA dynamic state: confined, anterograde vs retrograde for *Camk2a*, *Beta actin* and *Psd95.* n=59 cells. **/**** p<.01/.0001 Holm-Sidak’s multiple comparison test. **D.** Quantification of distance traveled for *Camk2a*, *Beta actin* and *Psd95.* n = 59 cells. **E.** Quantification of mRNA velocity for anterograde and retrograde for *Camk2a (1.013 +/− 0.450; −1.083 +/− .480)*, *Beta actin (1.054+/− 0.493; −1.110 +/− 0.452)* and *Psd95 (0.997 +/− 0.426; - 1.111 +/− 0.479).* n = 59 cells.

### Translational inhibition alters mRNA dynamics within the dendrite

With the above measurements of basal mRNAs dynamics, we next assessed if we could alter their dynamic properties. We first assessed if perturbing the translational status of an mRNA could affect its motility. To alter the translation status of an mRNA we used two mechanistically distinct translational elongation inhibitors (Figure 3A): puromycin, which causes release of the nascent peptide chain and ribosomal dissociation from the mRNA_35_, and anisomycin, which freezes elongating ribosomes on mRNAs_36_. As such, puromycin promote the transition to a ribosome-free mRNA state, whereas anisomycin causes ribosome accumulation on the mRNA. Using our analysis pipeline, we quantified the effects of these treatments on mRNA dynamics (Figure 3B-E). For *Camk2α* and *Beta actin*, puromycin displacement of ribosomes led to enhanced motility (reduced confinement) (*Camk2a*: 0.59 +/− 0.10 vs 0.43 +/− 0.13; *Beta actin*: 0.56 +/− 0.13 vs 0.44 +/− 0.12. Mean +/− SEM) (Fig 3B) and distance traveled (*Camk2a*: 18.14+/− 20.26 vs 24.48 +/− 25.70; *Beta actin*: 19.64 +/− 20.73 vs 27.59 +/− 24.78) (Fig 3C). In contrast, *Psd95* motility (*Psd95*: 0.51 +/− 0.14 vs 0.45 +/− 0.14) (Figure 3B) and distance traveled (Figure 3C) (*Psd95*: 20.71+/− 19.21 vs 17.44 +/− 20.41) was not significantly changed by puromycin treatment. This difference may reflect a higher basal translational state for *Camk2α* and *Beta actin* mRNAs, consistent with *Psd95* mRNA being slightly more dynamic relative to the other two mRNAs (Figure 2). Since displacing ribosomes for *Camk2α* and *Beta actin* enhanced mRNA motility-we predicted that freezing the ribosomes on the mRNA should lead to the opposite effect. Indeed, anisomycin treatment led to decreased mRNA motility (Figure 3D) for all three mRNAs (*Camk2a*: 0.54 +/− 0.10 vs 0.64 +/− 0.11; *Beta actin*: 0.55 +/− .09 vs 0.63 +/− .09; *Psd95*: 0.49 +/− 0.09 vs 0.58 +/− .09. Mean +/− SEM) and distance traveled (Figure 3E) for *Beta actin* and *Psd95* (*Camk2a*: 21.39 +/− 19.24 vs 20.06 +/− 17.85; *Beta actin*: 25.65 +/− 22.24 vs 22.32 +/− 17.66; *Psd95*: 21.47 +/− 17.70 vs 16.96 +/− 16.75). Interestingly, neither displacing nor freezing ribosomes had an effect on the active transport velocity of any mRNA (Supplemental Figure 2B&C). Furthermore, neither puromycin nor anisomycin affected the stationary population (Supplemental figure 2D&E). Given that transport was unaffected by either perturbation of translation, (Supplemental Figure 2B&C), our data is consistent with the idea that mRNAs are transported in a quiescent non-translating state_23,37_. Taken together, these data illustrate that the translational status of a given mRNA will affect its dynamics within the dendrite.

**Figure 3:**
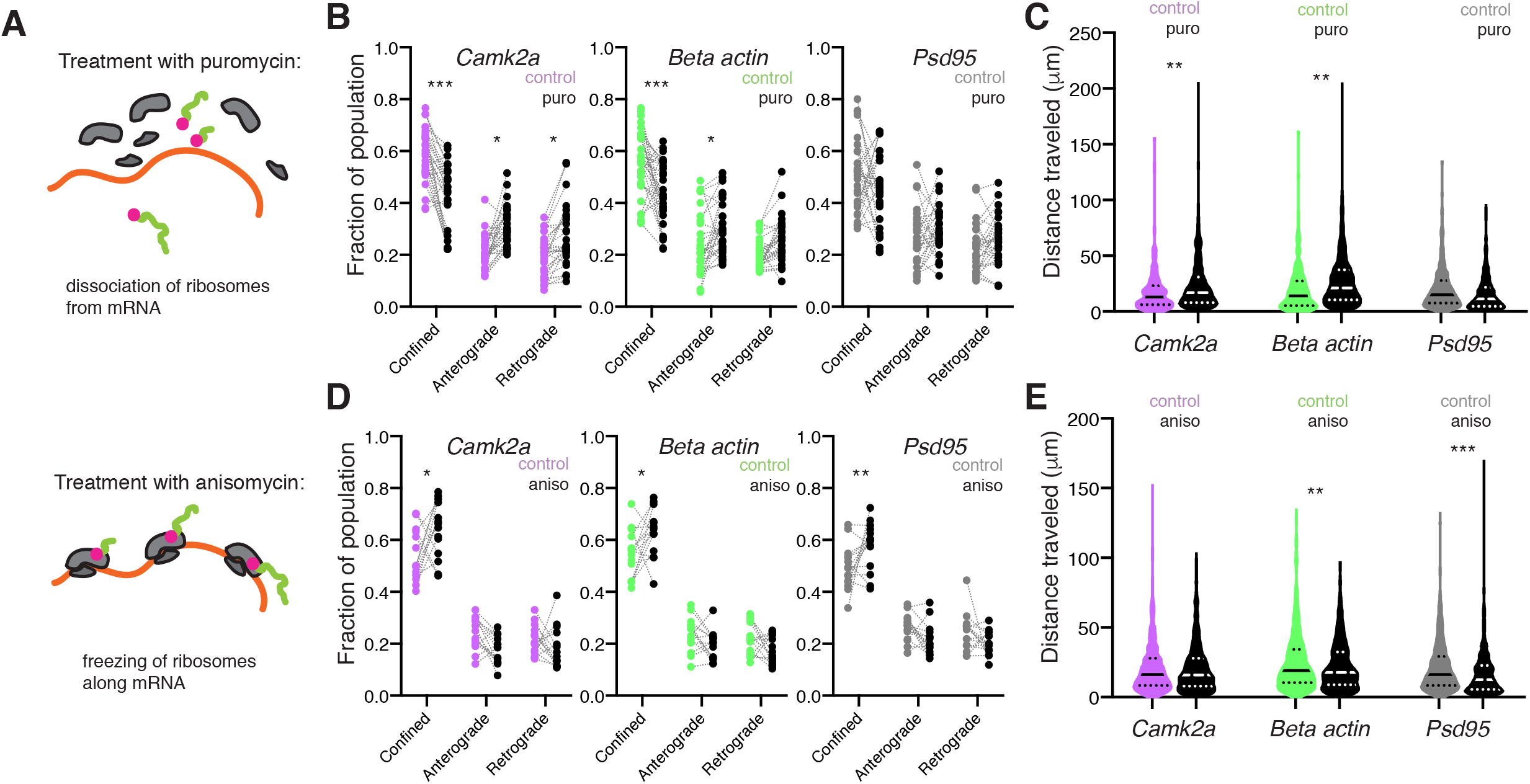
Manipulating ribosome association of mRNAs results in transcript specific alterations in mRNA dynamics. **A.** Schematic representation of the effects of puromycin (right) or anisomycin (left) on ribosomal association with mRNA. Puromycin results in ribosomal subunit disassembly whereas Anisomycin results in stalling of elongating ribosomes. **B.** Quantification of mRNA dynamic state: confined, anterograde vs retrograde for *Camk2a*, *Beta actin* and *Psd95* for control vs puro treated samples. n=29 cells per condition. */*** p<.05/.001. Sidak’s multiple comparisons test. **C.** Quantification of distance traveled for *Camk2a*, *Beta actin* and *Psd95* for control vs puro treated samples. N = 29 cells per condition. ** p<.01 Sidak’s multiple comparisons test. **D.** Quantification of mRNA dynamic state: confined, anterograde vs retrograde for *Camk2a*, *Beta actin* and *Psd95* for control vs aniso treated samples. n = 15 cells per condition. */** p<.05/.01 Sidak’s multiple comparisons test. **E.** Quantification of distance traveled for *Camk2a*, *Beta actin* and *Psd95* for control vs aniso treated samples. n=15 cells per condition. **/*** p<.01/.001 Sidak’s multiple comparisons test.

### Plasticity stalls mRNA transport and accumulates mRNAs near dendritic spines

We next assessed if we could modulate mRNA dynamics with physiologically relevant manipulations-specifically synaptic plasticity. We examined how mRNA dynamics are altered during two forms of protein synthesis-dependent plasticity, chemically-induced long-term potentiation (cLTP)_38_ and metabotropic glutamate receptor-mediated long-term depression (mGluR-LTD)_39_ (Supplemental Figure 3A&B). We induced cLTP (Figure 4A&B, Supplemental figure 3C) or mGluR-LTD (Figure 4C&D, Supplemental Figure 3D) and monitored mRNA dynamics immediately after induction. Induction of cLTP led to decreased mRNA motility (Figure 3D) for all three mRNAs (*Camk2a*: 0.54 +/− 0.09 vs 0.63 +/− 0.08; *Beta actin*: 0.53 +/− .10 vs 0.62. +/− .06; *Psd95*: 0.47 +/− 0.06 vs 0.58 +/− .10. Mean +/− SEM.) and reduced distance traveled (Figure 4B) (total distance traveled in microns: *Camk2a*: 23.36 +/− 19.86 vs 20.54 +/− 18.97; *Beta actin*: 18.95 +/− 19.75 vs 16.25 +/− 14.52; *Psd95*: 28.95 +/− 23.29 vs 22.87 +/− 18.91). Induction of mGluR-LTD led to decreased mRNA motility (Figure 4C) for all three mRNAs (*Camk2a*: 0.54 +/− 0.09 vs 0.64 +/− 0.09; *Beta actin*: 0.56 +/− .10 vs 0.63 +/− .10; *Psd95*: 0.48 +/− 0.06 vs 0.62 +/− .07. Mean +/− SEM) and reduced distance traveled (Figure 4D) (total distance traveled in microns: *Camk2a*: 28.81 +/− 20.55 vs 23.66 +/− 18.48; *Beta actin*: 23.37 +/−19.34 vs 17.97 +/− 14.00; *Psd95*: 25.54 +/− 20.68 vs 21.84 +/− 16.05). To summarize, we observed a significant decrease in the time all 3 mRNAs spent moving and a reduced distance traveled within the dendrite for both forms of plasticity.

**Figure 4:**
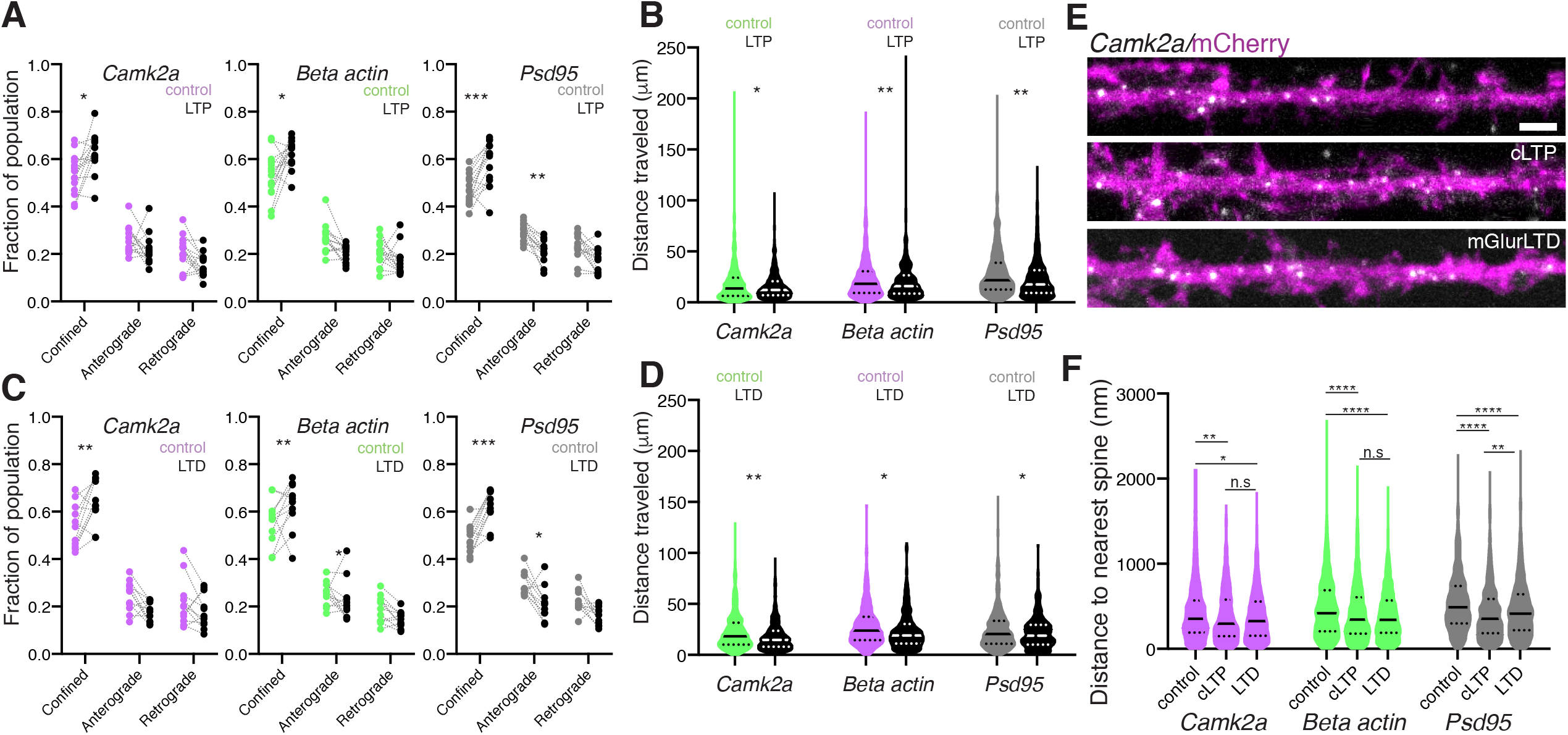
Two distinct forms of synaptic plasticity attenuate mRNA dynamics near dendritic spines. **A.** Quantification of mRNA dynamic state: confined, anterograde vs retrograde for *Camk2a*, *Beta actin* and *Psd95* for control vs cLTP induced samples. n=14 cells per condition. */*** p<.05/.001. Sidak’s multiple comparisons test. **B.** Quantification of distance traveled for *Camk2a*, *Beta actin* and *Psd95* for control vs cLTP induced samples. n=14 cells per condition. ** p<.01. Sidak’s multiple comparisons test. **C.** Quantification of mRNA velocity for anterograde and retrograde for *Camk2a*, *Beta actin* and *Psd95* for control vs cLTP induced samples. n = 14 cells per condition. **D.** Quantification of mRNA dynamic state: confined, anterograde vs retrograde for *Camk2a*, *Beta actin* and *Psd95* for control vs mGluR-LTD induced samples. n=14 cells per condition. */** p<.05/.01. Sidak’s multiple comparisons test. **E.** Quantification of distance traveled for *Camk2a*, *Beta actin* and *Psd95* for control vs mGluR-LTD induced samples. n = 14 cells per condition. **/*** p<.01/.001. Sidak’s multiple comparisons test. **F.** Example images of Camk2a RNA FISH signal in neurons volume filled with mCherry under control, +cLTP and +mGluR-LTD conditions. Scale bar = 2.5um. **G.** Quantification of mRNA distance to the nearest spine reveals a slight decrease in the distance for all three mRNAs during plasticity. Control, cLTP, mGluR-LTD +/− SD: *Camk2a* (411.6 +/− 300.7; 376.8 +/− 292.5; 375.0 +/− 275.4), *Beta actin* (474.3 +/− 327.1; 413.8 +/− 312.3; 391.0 +/− 275.8) and *Psd95* (537.2 +/− 334.3; 403.3 +/− 283.7; 454.6 +/− 304.7) */**/**** p<.05/.01/.0001; Dunn’s multiple comparisons test. n = 15 cells per condition.

To assess more precisely the location of mRNA deposition during these enhanced periods of mRNA confinement, we performed high resolution single molecule fluorescent *in situ* hybridization in dendrites immediately after induction of cLTP and mGluR-LTD (smFISH, see methods) (Figure 4E&F). We measured the mean distance of an mRNA granule to its nearest dendritic spine and found that this distance decreased significantly with both cLTP and mGluR-LTD induction for all three mRNAs (Figure 4F). Taken together with the altered dynamics observed with the molecular beacons (Figure 4A&C) our data support increased spine association of these mRNAs during plasticity. This enhanced association may fuel local translation of these mRNAs to induce and maintain both forms of structural plasticity.

### Exploring the dynamics of protein synthesis in real-time

To assess directly whether translation of these three mRNAs is enhanced during cLTP and mGluR-LTD, we used translational reporters_18_ (Figure 5) comprising an optimized super-folder GFP_40_ (sfGFP, see methods) flanked by the corresponding dendritically enriched 3’UTR_41_ of *Cam2a*, *Beta actin*, or *Psd-95*. The 3’UTR was included to confer both transcript-specific localization and translational regulation to the translational reporter_41_. We used cell-wide fluorescence recovery after photobleaching (FRAP) to visualize newly synthesized proteins. Following whole cell photobleaching, we measured the emergence and time course of the protein synthesis-dependent fluorescence signal to assess the kinetics and extent of the translational responses for each mRNA. We found that all three transcript-specific reporters showed protein synthesis-dependence in their recovery compared to the no UTR control (Figure 5D, Supplemental Figure 4, black vs gray curves), indicating that these reporters are effective readouts for active translation. The induction of either cLTP or mGluR-LTD resulted in an enhancement of the mobile fraction (Figure 5D) for both the CAMK2α and BETA ACTIN reporter and total fluorescence recovery (Supplemental Figure 4) indicating enhanced synthesis. Conversely, PSD-95 showed enhanced translation following cLTP induction (Figure 5D, Supplemental Figure 4), but no change following mGluR-LTD. Interestingly, we noticed a strong bias for the emergence of fluorescence in spines fluorescence during cLTP and mGluR-LTD (Figure 5C). To assess if this represented a bias for spine accumulation of newly synthesized protein, we assessed the ratio of fluorescence recovery for the spine over the shaft (Figure 5E). Plasticity induction indeed resulted in a higher rate of spine recovery, except for PSD-95 during mGluR-LTD, suggesting that local protein production near spines fuels the emergence of new fluorescence signal. Taken together, these data indicate that there may be a functional disconnect between transcript-specific changes in mRNA dynamics seen during plasticity (Figure 3) and the downstream translational state of that mRNA species.

**Figure 5:**
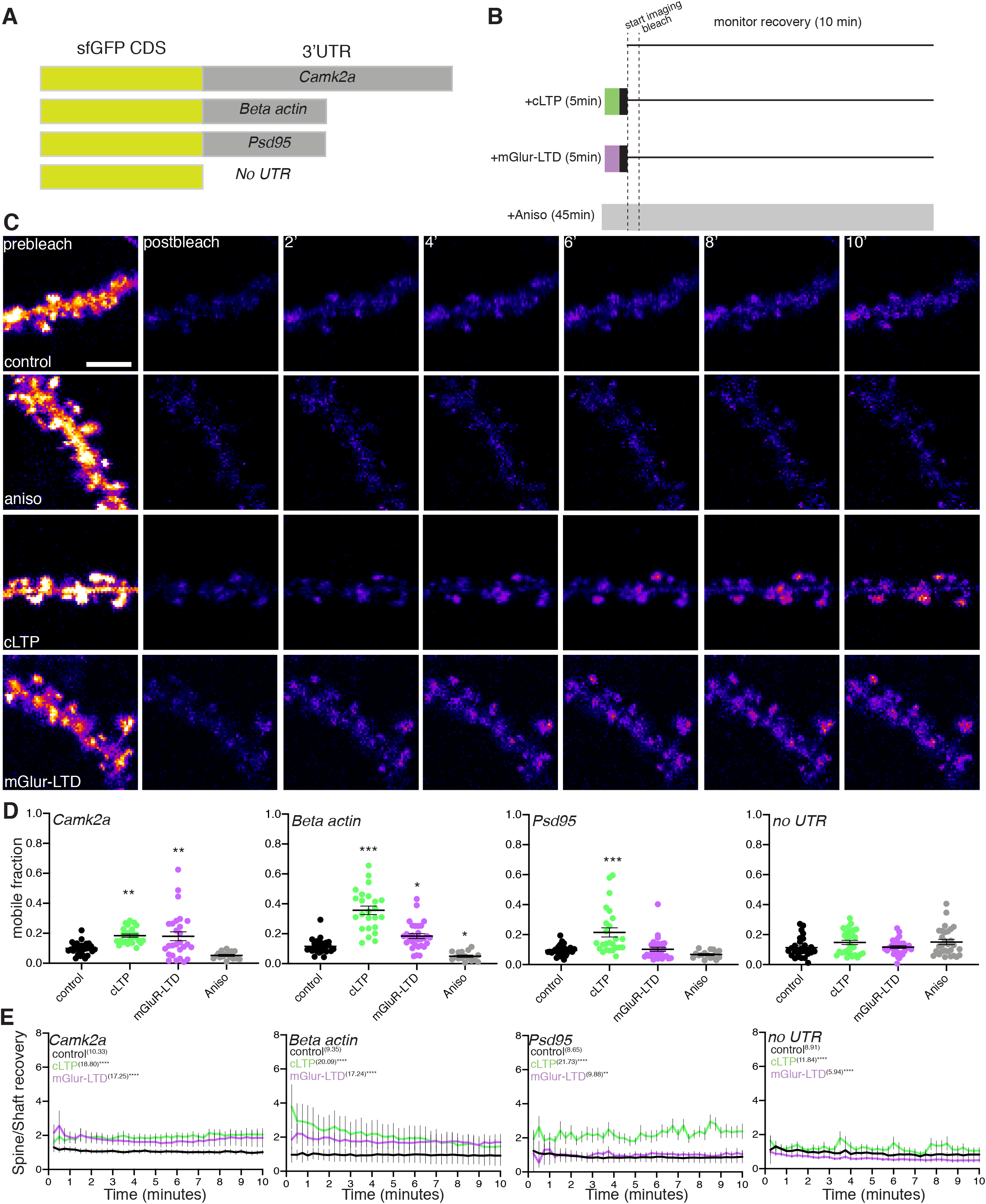
3’UTR regulated translation during synaptic potentiation and depression. **A.** Scheme for the reporters used to assess real-time 3’UTR regulated translational dynamics in live neurons. **B.** Scheme of workflow for the treatment and visualization of plasticity regulated protein synthesis. Following induction of cLTP or mGluR-LTD, the indicated pharmacological treatment was washed out (black box) and the samples were then imaged for 2 minutes every 15 seconds to acquire a baseline measurement. The cells were then bleached- and the fluorescence recovery was monitored every 15 seconds for 10 minutes. For controls, either no treatment or treatment with the translational inhibitor anisomycin was used for comparison. **C.** Example image of a Camk2a sfGFP reporter under control (top) +aniso (second row) +cLTP (third row) or +mGluR-LTD conditions before bleaching and during the phase of fluorescence recovery. Scale bar = 5um. **D.** Mobile fractions calculated for the translational reporters during control, plasticity induction and anisomycin treatment. *\*/**/\*\*\** p <.05/.01/.001. Dunnett’s multiple comparisons test for treated vs control condition for each construct; n = >14 cells per condition. **E.** Recovery rate of fluorescence of dendritic spines to shafts demonstrates a bias for spine fluorescence recovery during plasticity. Kruskal-Wallis test. *\*/**/***/\*\*\*\** p <.05/.01/.001/.0001. n= >14 cells per condition.

To assess if the same pattern of translation following plasticity was also observed with endogenous transcripts, we used CRISPR/Cas9 gene editing in neurons_42_ to tag endogenous CAMK2α or BETA ACTIN (n-terminal) or PSD-95 (c-terminal) protein with the fast-folding Venus fluorescent protein_43_ (Figure 6A, Supplemental Figure 4). Similar to sfGFP, the fast-folding nature of Venus (t_1/2 maturation_ = 2-5 minutes) allowed us to rapidly assess the translational regulation of these three proteins. All three proteins were successfully tagged and exhibited their characteristic localization patterns (Supplemental figure 5). Venus-tagged CAMK2α and BETA ACTIN were enriched in axons and dendrites, and most strongly enriched in spines and boutons; while PSD-95 tagged Venus was exclusively enriched within postsynaptic compartments. As before, we performed cell-wide fluorescence recovery after photobleaching (FRAP, see methods) (Figure 6B-E; Supplemental Figure 6A-D, Supplemental Table 2). We tracked the emergence and time course of the protein synthesis-dependent fluorescence signal to assess the kinetics and the extent of the translational responses for each mRNA (Supplemental Figure 6B-D). For all proteins, treatment with the protein synthesis inhibitor anisomycin significantly reduced the emergence of new fluorescent signal (Supplemental Figure 6B) and attenuated the mobile population during recovery (Figure 6C) (CAMK2α: 0.21 +/− 0.01 vs 0.13 +/− 0.06; BETA ACTIN: 0.21 +/− 0.01 vs 0.11 +/− 0.007; PSD-95: 0.35 +/− 0.03 vs 0.21 +/− 0.02 Mean +/− SEM) indicating that we were able to visualize protein synthesis in real-time for endogenous proteins. Similar to what we observed with the translation reporters, we found that cLTP induction significantly enhanced the emergence of fluorescence for all three proteins (Supplemental Figure 6C) and enhanced the mobile population (Figure 6D) (CAMK2α: 0.23 +/− 0.02 vs 0.3 +/− 0.01; BETA ACTIN: 0.21 +/− 0.01 vs 0.30 +/− 0.02; PSD-95: 0.35 +/− 0.03 vs 0.45 +/− 0.02 Mean +/− SEM), indicating enhanced protein synthesis. Induction of mGluR-LTD, however, elicited a transcript-specific enhancement of protein synthesis for CAMK2α and BETA ACTIN but not PSD-95 (CAMK2α: .20 +/− 0.02 vs 0.29 +/− 0.02; BETA ACTIN: 0.23 +/− 0.02 vs 0.32 +/− 0.02; PSD-95: 0.33 +/− 0.03 vs 0.37 +/− 0.02. Mean +/− SEM), (Figure 6E, Supplemental Figure 6D).

**Figure 6:**
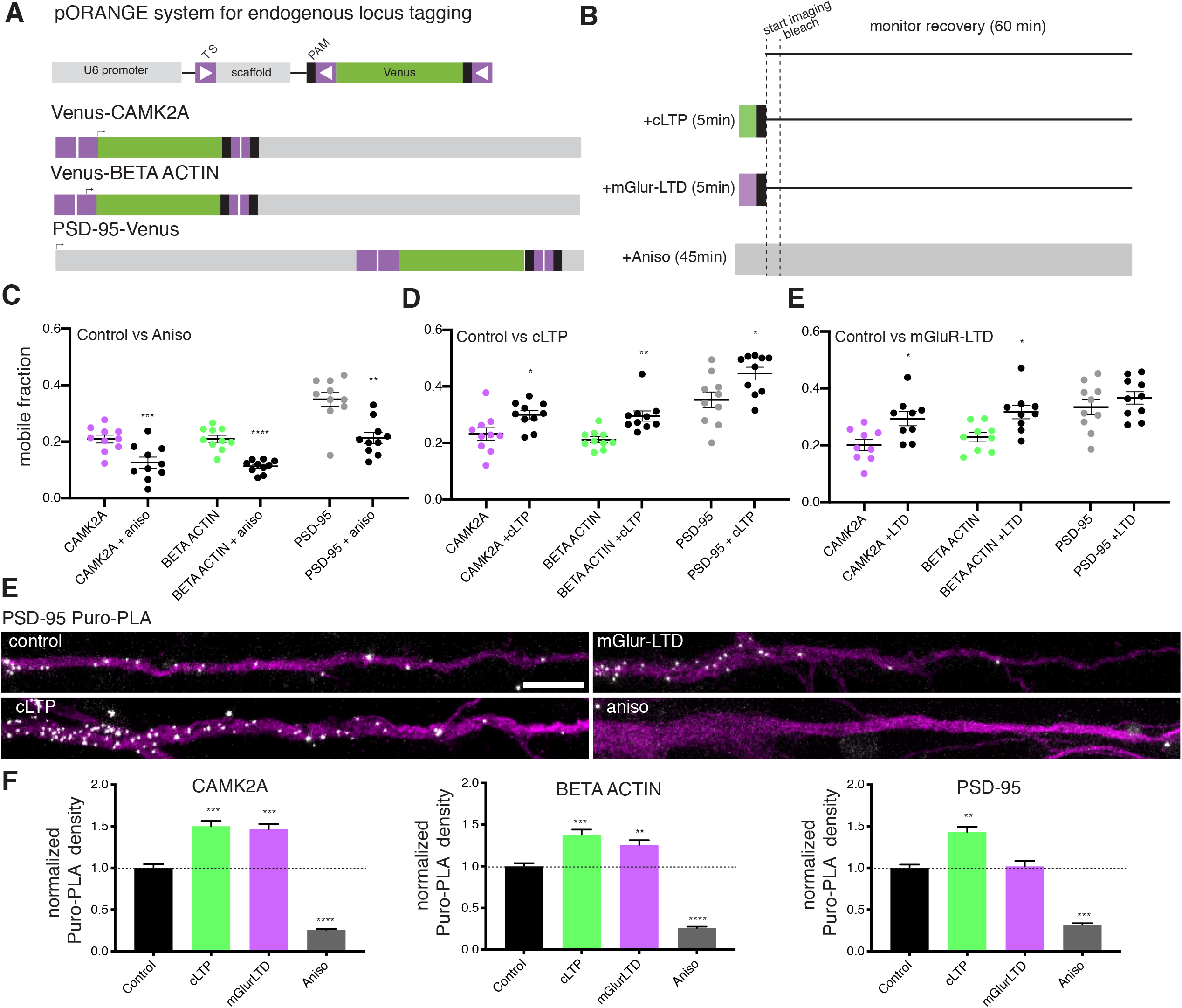
Visualizing endogenous protein translation in real time during synaptic potentiation and depression. **A.** Scheme for the modification of endogenous gene tagging of Camk2a/Beta actin/Psd95 with Venus fluorescent protein. T.S = targeting sequence, arrow indicates start codon. **B.** Scheme of workflow for the treatment and visualization of plasticity regulated protein synthesis. Following induction of cLTP or mGluR-LTD, the pharmacological treatment was washed out (black box) and the samples were then imaged for 2 minutes every 15 seconds to acquire a baseline measurement. The cells were then bleached- and the fluorescence recovery was then monitored every 15 seconds for 60 minutes. For controls, either no treatment or treatment with the protein synthesis inhibitor anisomycin was used for comparison. **C.** Mobile population during the FRAP recovery for Venus-CAMK2a, Venus-BETA ACTIN and PSD-95-Venus during control and anisomycin treatment. n = 10 cells per condition. Two-tailed paired T-test. **/***/**** p<.01/.001/.0001. **D.** Mobile population during the FRAP recovery for Venus-CAMK2a, Venus-BETA ACTIN and PSD-95-Venus during control and cLTP. n = 10 cells per condition. Two-tailed paired T-test. */** p<.05/.01. **E.** Mobile population during the FRAP recovery for Venus-CAMK2a, Venus-BETA ACTIN and PSD-95-Venus during control and mGluR-LTD. n = 9-10 cells per condition. Two-tailed paired T-test. * p<.05. **F.** Example images dendritically localized (MAP2, magenta) Puro-PLA signal for PSD-95 (white) under control and stimulated conditions. Scale bar = 10um. **G.** Puro-PLA quantification reveals local protein synthesis underlies the translational responses of CAMK2a, BETA ACTIN and PSD-95 during cLTP and mGluR-LTD. n = 85 cells per condition. Dunnett’s multiple comparisons test. */**/*** p<.05/.01/.001.

To obtain better temporal and spatial resolution on the translational responses to plasticity induction, we validated the above data with a method that couples general metabolic labeling of nascent proteins with a specific label for a protein-of-interest using the proximity ligation assay (Puro-PLA_44_, see methods, Supplemental Figure 7A). With this method we again examined changes in local dendritic protein synthesis during plasticity (Figure 6F&G, Supplemental Figure 7B&C). Consistent with previous observations_44_, we observed that there is a protein synthesis dependence to the Puro-PLA signal for all 3 proteins (black vs grey bars) (CAMK2α: 0.26 +/− 0.13; BETA ACTIN: 0.26 +/− 0.14; PSD-95: 0.32 +/− 0.17. Mean +/− SEM). Furthermore, cLTP induction enhanced the dendritic synthesis of all three mRNAs (CAMK2α: 1.50 +/− 0.58; BETA ACTIN: 1.38 +/− 0.57; PSD-95: 1.43 +/− 0.58. Mean +/− SEM); while induction of mGluR-LTD only enhanced the synthesis of CAMK2α and BETA ACTIN (CAMK2α: 1.47 +/− 0.55; BETA ACTIN: 1.26 +/− 0.52; PSD-95: 1.02 +/− .61. Mean +/− SEM). These data not only validate and support our real time translational response observations, but also support that this translation can occur, at least in part, locally within the dendrite where we detect changes in mRNA dynamics. Taken together, these data indicate that alterations in mRNA dynamics and protein synthesis underlie the manifestation of specific forms of synaptic plasticity.

## Discussion

Here we investigated the interplay between mRNA dynamics and translation within neuronal dendrites during two different forms of synaptic plasticity. We characterized the dynamics and translation of three individual endogenous mRNAs: *Camk2a*, *Beta actin* and *Psd95*, during basal neuronal activity and plasticity. In live hippocampal neurons we provide evidence that mRNAs exist in heterogenous copy number organizational states (Figure 1). The preference for single mRNA copy vs higher order state appears to be a transcript-specific feature, as *Camk2a* and *Psd95* exhibit an enhanced preference for higher order multimeric states compared to the previously described single mRNA state of *Beta actin*_29_. We found that ribosome association directly influences mRNA dynamics-suggesting that mRNA translation is likely restricted to non-transporting mRNAs exclusively within the dendrite. Consistent with our data, previous work in fibroblasts_45_ and axons_32_ has shown that diffusion of *Beta actin* increases when ribosomes are displaced. These data fit with recent proteomic analysis on isolated mRNA transport granules_46_ which detected only a subset of ribosomal proteins associated with granules.

In our experiments, mRNAs were persistently sequestered, exhibiting reduced mobility, following plasticity (e.g. for entire duration of imaging, ~20 min). The mechanisms underlying the initial capture and enduring sequestration of mRNAs are not well understood. Previous work with *Beta actin* capture during glutamate uncaging suggest that it involves actin remodeling_22_. Indeed, the structural spine plasticity characteristic of both cLTP and mGluR-LTD involve modulation of actin cytoskeleton_47_. Whether actin remodeling broadly promotes mRNA sequestration at or near dendritic spines remains assessed. The translation of all three mRNAs was enhanced by LTP, but LTD only enhanced the translation of *Camk2a* and *Beta actin*. *Psd95* mRNA has previously been characterized as an mGluR-LTD regulated transcript_16,48,49_. However, following washout of the mGluR agonist (similar to the conditions used here) PSD-95 protein is rapidly degraded_50_. The above literature is consistent with our data, where *Psd95* mRNA is retained at spines but its protein synthesis is unchanged relative to the control state. Taken together, our data dissociates the accumulation of mRNAs near synapses from their translational status_22_. Given specific signaling cascades are turned on by distinct forms of plasticity_2_, these cascades likely influence changes in post-translational modifications of RNA binding proteins on particular transcripts-regulating their translatability. Consistent with this, activation of PKA signaling alone is sufficient to enhance dendritic protein synthesis_11_. LTP, known classically for its dependence on CamkII signaling_51_ also triggers a number of other classical signaling cascades including PKA_52_, PKC_53,54_, MAPK/ERK_55_, PI3K_56_, mTOR_57_ and Src_58_. mGluR-LTD on the other hand, is less clear in its signaling requirements –it involves activation of PKC_59_ and PI3K/AKT/mTOR_60_ however the role of CamkII_61,62_is debated. Once sequestered by an active synapse, mRNAs could be “synaptically decoded” by the activation of these signaling pathways to determine whether and when a given mRNA species will be translated or not.

## Acknowledgements

We thank I. Bartnik, N. Fuerst, A. Staab, D. Vogel and C. Thum for the preparation of cultured neurons.

## Author contributions

P.G.D.A. designed, conducted and analyzed experiments, and wrote the paper. C.P designed, conducted and analyzed experiments. R.K. designed and conducted experiments. A.H designed experiments and supervised the project. E.M.S. designed experiments, supervised the project, and wrote the paper. All authors edited the paper.

## Data and materials availability

All data is available in the manuscript or the Supplemental materials.

## Methods

### Molecular Beacon Structure and Design

#### Solid-phase synthesis

Milli-Q water was treated with DEPC (0.1%) overnight and autoclaved.

The following oligonucleotides were synthesized on an *ABI392* instrument:

PSD95_1: 5’-M CACGACCAUCCCUCCCCUUUUCCCAAAAAAAUAUCGUG Q_1_ - 3’
PSD95_2: 5’-M CACGAAUAAAAUCCCAGAAAAAAAAAAAGCCUCGUG Q_2_ S −3’
CAMK2_1: 5’- M CACGAGGUAAAAACUUCCCCUCACUCCUCUUCCUCGUG Q_1_-3’
CAMK2_2: 5’-M CACGAUUUUUCUUCUUUUUUGUUUUGCUCUUCGUG Q_2_ S −3’
Beta Actin_1:5’-M CACGACAAAACAAAACAAAAAAACUUAAAAAAAUCGUG Q_1_ −3’
Beta Actin_2:5’-M CACGAUUCACCGUUCCAGUU UUUAAAUCCUGUCGUG Q_2_ S - 3’
M = Fmoc-Amino-DMT C-3 CED phosphoramidite (*ChemGenes*)
Q_1_ = 3’-BHQ-2 CPG 1000 (*LinkTech*)
Q_2_ = BBQ-650®-(DMT)-CE-Phosphoramidite (*LinkTech*)
S = 3’-Spacer C3 SynBase™ CPG 1000/110 (*LinkTech*)
A = 2’-OMe-Pac-A-CE Phosphoramidite (*LinkTech*)
C = 2’-OMe-Ac-C-CE Phosphoramidite (*Linktech*)
G = 2’-OMe-*i*Pr-Pac-G-CE Phosphoramidite (*LinkTech*) U = 2’-OMe-U-CE Phosphoramidite (*LinkTech*)

For all synthesized oligonucleotides, Pac_2_O was used as capping reagent. 0.3 M BTT (*emp Biotech*) was used as activator. Coupling time for A, C, G, U and M was 6 minutes, for Q_2_ 15 minutes. Synthesis was performed in DMTr-On mode. The cyanoethyl groups were removed by flushing the columns with 20% diethylamine (*emp Biotech*) for 10 minutes, followed by washing with MeCN, Argon and drying in vacuum. Cleavage from the solid-phase was performed with aqueous ammonia (32%) (*Merck*) for 4 hours at room temperature. After spin filtration, the solvent was removed at 4 °C using a vacuum concentrator (*SpeedVac™*, *Thermo Fischer*).

### Purification

The DMTr-On oligonucleotides were purified by RP-HPLC on an *Agilent 1200* equipped with a *waters XBridge BEH C18 OBD* column (300 Å, 5 μm, 19×250 mm, 4 mL/min, 60 °C). As solvents 400 mM hexafluoroisopropanol (*fluorochem*), 16.3 mM Et_3_N (*Merck*), pH 8.3 and MeOH (*Fluka*) were used with a gradient from 5% to 100% MeOH in 30 minutes. After separation, the solvent was evaporated in a vacuum concentrator at 4 °C.

The DMTr group was removed by incubation of the oligonucleotides in 400 μL 80% aqueous AcOH (*Merck*) at room temperature for 20 minutes, followed by evaporating the solvent in a vacuum concentrator at 4 °C. The RNAs were again purified by RP-HPLC under the same conditions as above.

### Fluorophore labeling

10 nmol of each RNA were dissolved in 150 μL borate-buffer (0.1 M sodium tetraborate (*Merck*), pH 8.4). PSD95_1, CAMK2_1 and Beta Actin_1 were incubated with 200 nmol ATTO565 NHS (*ATTO-TEC*), dissolved in 50 μL DMF (*lumiprobe*, labeling grade), for 4 hours at 37 °C. PSD95_2, CAMK2_2 and Beta Actin_2 were incubated with 200 nmol ATTO647N NHS (*ATTO-TEC*), dissolved in 50 μL DMF, for 4 hours at 37 °C. Buffer and the excess of fluorophore were removed by size exclusion chromatography (*NAP 25*, *GE Healthcare*). The solvent was evaporated at 4 °C using a vacuum concentrator. The residue was purified by RP-HPLC on an *Agilent 1200* equipped with an *Xbridge BEH C18 OBD* (300 Å, 3.5 μm, 4.6×250 mm, 1 mL/min, 60 °C). As solvents 400 mM hexafluoroisopropanol, 16.3 mM Et_3_N, pH 8.3 and MeOH were used with a gradient from 5% MeOH to 100% MeOH in 50 minutes.

### Sample preparation for in vivo use

For use in living cells the remaining HPLC buffer ions had to be removed. Therefore, the oligonucleotides were dissolved in 0.3 M NaOAc (*Merck*) (10 μL per 1 nmol RNA). EtOH (*Fluka*, prechilled to −20 °C, 40 μL per 1 nmol RNA) was added. The mixture was cooled to −20 °C for at least 6 hours. The precipitant was pelletized by centrifugation at 4 °C, 20000 g for 20 minutes. The residue was redissolved in 0.3 M NaOAc and the precipitation steps were repeated 3 times. To remove sodium ions, the oligonucleotides were desalted using a 1k cut-off membrane filter (*Microsep Advance Centrifugal Devices with Omega Membrane 1K*, *PALL*). Before adding the oligonucleotides, each filter was washed 5 times with DEPC water at 15000 g, 15 °C for 20 minutes. The desalting step was repeated 3 times.

### Characterization

Analytical RP-HPLC was performed on an *Agilent 1200* equipped with a *BEH C18 OBD* (300 Å, 3.5 μm, 4.6×250 mm, 1 mL/min, 60 °C). As solvents 400 mM hexafluoroisopropanol (*fluorochem*), 16.3 mM Et_3_N (*Merck*), pH 8.3 and MeOH (*Fluka*) were used with a gradient from 5% to 100% MeOH in 39 minutes. ESI-MS spectra were recorded on a *Bruker micrOTOF-Q* device in negative ionization mode.

### Hippocampal neurons

Dissociated rat hippocampal neuron cultures were prepared and maintained as described previously_1_. Cells were plated at a density of 30 − 40 × 10_3_ cells/cm² on poly-D-lysine coated glass-bottom Petri dishes (MatTek). Hippocampal neurons were maintained and matured in a humidified atmosphere at 37°C and 5% CO2 in growth medium (Neurobasal-A supplemented with B27 and GlutaMAX-I, life technologies) for 18-21 days *in vitro* (DIV) in vitro to ensure synapse maturation. All experiments complied with national animal care guidelines and the guidelines issued by the Max Planck Society and were approved by local authorities.

### Transfection of plasmid DNA

For transfection of fluorescent proteins and reporters, DIV17-19 neurons were transfected with mCherry-C1 (clonetech), myr-sfGFP translational reporters (described below) or myr-Venus using Effectene (Qiagen), as previously described_2_. pCAG:myr-Venus_3_ was a gift from Anna-Katerina Hadjantonakis (Addgene plasmid # 32602; http://n2t.net/addgene:32602; RRID:Addgene_32602). Transfected cells were imaged or fixed (described below) 12-18 hours post transfection.

### Transfection of molecular beacons

For transfection of molecular beacons, DIV17-19 neurons were transfected with Attractene (Qiagen). For each Mattek dish, 20pmol of molecular beacon was resuspended in 75uL of buffer EC (Qiagen) along with 2uL of Attractene. The beacon-attractene mix was incubated for 20 minutes at room temperature before being added to neurons. Samples were imaged 1-12 hours post transfection. Prior to imaging, samples were washed in fresh media to remove non-transfected beacons.

### Electroporation of plasmid DNA

Following isolation, 1 million hippocampal neurons were spun down at 500rpm for 5 minutes at 4C°. Cells were resuspended in electroporation solution (Lonza) along with 1ug pORANGE plasmid DNA construct. Cells were electroporated with the hippocampal/cortical high viability protocol (Lonza) and resuspended in 2mL cell growth media. Cells were then plated at a density of 100×10_3_ in Mattek dishes coated with poly-D lysine for 2 hours to allow for cell attachment. Following attachment, 1.3mL media was added, and cells were fed with 500uL fresh neuronal growth media once a week until the time of experiments.

### Cell treatments

Drugs treatments were performed as follows: For puromycin labeling experiments (Puro-PLA), cultured neurons were treated with 10 μM puromycin for 5-10 min. For Puro inhibition experiments, cultured neurons were treated with 100 μM puromycin for 5 minutes. Anisomycin treatment (40 μM) was performed 20-45 min prior to puromycin labeling, FRAP or molecular beacon experiments, and was kept in the media through the duration of experiment. mGluR-LTD was induced using (S)-3,5-Dihydroxyphenylglycine hydrate (DHPG; 100 μM) for 5 min and then washed out. cLTP was induced as previously described_2_ in E4 buffer supplemented with B27, Glutamax and MEM amino acids (Thermofisher). The day before the experiment 50uM APV (Tocris) was added to neuronal cultures. The day of induction, neurons were incubated in Mg2+ free E4 media supplemented with 200uM glycine (sigma) and 100uM picrotoxin (Tocris) for 5 minutes. Following induction cells were washed and returned to normal media or E4 with calcium and magnesium.

### Imaging of molecular beacons

Investigation of the mRNA dynamics was carried out using a Leica DMi8 TIRF microscope. Differential interference contrast (DIC) microscopy was used to identify neurons with well-isolated dendrites. mRNA dynamics were recorded for 20 minutes at a rate of 1 Hz in epi-fluorescence mode. ATTO565 fluorophores were excited using a 561 nm diode laser which provided 1.8 kW/cm_2_ of intensity at the sample plane. ATTO647n was imaged with a 638 nm laser which produced 2.0 kW/cm_2_ at the sample plane. The fluorescence was recorded with a scientific-CMOS camera (Leica-DFC9000GT). The exposure time was fixed to 200 ms and 2×2 camera binning and set the digitalization to 12 bit (low noise) was used to limit the data volume. A 100x oil objective (HC PL APO 100x/1.47 OIL) was used to record a field-of-view of 133 μm x 133 μm. With these settings our pixel size was 130 nm, matching the Nyquist sampling frequency. Neurons were left in their glia-conditioned neurobasal, B27 and glutamax media owing to a Pecon TempController 2000-1 and a Pecon CO_2_-Controller 2000 which kept the samples at 37°C in a 5% CO_2_ atmosphere.

### Quantification of beacon number per puncta

To quantify the copy numbers of mRNAs within individual mRNA granules, a commercially synthesized oligo containing a single ATTO647n fluorophore (GATTA-Brightness R1 in 0.5 TBE and 11 mM Mg on glass slide, GattaQuant GmbH) was used as a normalization standard. This sample was imaged using the same settings for the molecular beacons at different laser powers, ranging from 90 μW (1%) to 6.85 mW (50%), at which point we could observe saturation of the fluorescence. To benchmark the mean counts recorded from a single ATTO647n fluorophore the maximum intensity around the detected puncta was measured and subtracted the neighboring background. This benchmark intensity was then set to the value n= 1 fluorophore and used to normalize the background-subtracted intensity recorded from hippocampal neurons.

### Quantifying mRNA dynamics

To extract information on the mRNA dynamics a custom MATLAB script was used. For each neuron a single dendrite was segmented by manually drawing its profile. The script was divided into filtering the images and rendering the puncta, tracking the mRNAs and extracting information regarding their dynamics. To render the puncta the background was subtracted by applying a mean filter. The pixel that represents the local maximum around a region of approximately 400 nm x 400 nm was then identified and selected. Puncta were rendered in a binary array. This pipeline was repeated each frame of the time series and exported as a movie. mRNA tracking was performed using the Motion-Based Multiple Object Tracking function of MATLAB taking the first 100 frames as training for the model. Any particle that did not appear in consecutive frames was discarded. After tracking was complete puncta that appear for longer than 20 frames (20 s) were retained and information such as the puncta coordinates, their distance travelled, their velocities and directionalities was extracted. Puncta that moved less than 500 nm throughout the imaging session were classified as fully stationary and were not included in directionality calculations. From the velocity data sets per puncta, the percent of time spent in the confined state was calculated by assessing the total number of frames a puncta was detected and how many frames this puncta exhibited a velocity from −500nm/s to 500nm/s. Similarly, the percent time spent anterograde or retrograde was calculated by the fraction of time >500nm/s or <− 500nm/s over the total number of frames detected. For all events detected in the cell, the average time spent in confined, anterograde or retrograde was calculated.

### Translational inhibitors and mRNA dynamics experiments

Beacons were transfected and imaged as described above, and puromycin and anisomycin were used at the concentrations indicated above. For assessing translational inhibitor effect on mRNA dynamics, two similarly looking neurons (containing a similar number of beacons and a similar morphology) were selected per Mattek dish. The first neuron was imaged as a control reference cell. Following the 20 minute imaging window for the control neuron, puromycin was added for 5 minutes or anisomycin was added for 20 minutes, prior to the start of imaging for the second, treated, cell. Pairwise assessment (Figure 3B&D) between the control and treated cell per dish was used to assess the effect of the drugs on mRNA dynamic properties.

### Spine size experiments

Neurons DIV17+ were transfected with myrVenus 12 hours prior to imaging and then imaged using a Leica DMi8 TIRF microscope. A 100x oil objective (HC PL APO 100x/1.47 OIL) was used to record a field-of-view of 133 μm x 133 μm using a 488 laser line. Samples were imaged for 10 minutes at baseline 1 frame every minute. Mock treatment, mGluR-LTD or cLTP was induced for 5 minutes, and samples were imaged 1 frame/minute during the induction phase. Following washout of drugs, neurons were imaged for 90 minutes post induction. For anisomycin treatments, anisomycin was added 20 minutes prior to the start of the experiment and kept in the media continuously throughout the experiment. Drift was corrected using the built-in correct 3D drift plugin in ImageJ/FIJI. An area of 5-10 spines per dendrite were then quantified over the imaging window, using the mean size of the first 10 baseline frames for normalization.

### Plasticity and mRNA dynamics experiments

Beacons were transfected and imaged and plasticity was induced as described above. For assessing the effect of plasticity on mRNA dynamics, two similar neurons (beacon number and morphology) were selected per mattek dish. The first neuron was imaged as a control reference cell. Following the 20 minute imaging window for the control cell, plasticity was induced and the stimulation washed out, prior to commencing imaging of the second, treated, cell. Pairwise assessment (Figure 4A&C) between the control and treated cell per dish was used to assess the effect of plasticity on mRNA dynamic properties.

### *Fluorescence In situ* hybridization

All steps were performed at room temperature, unless stated otherwise. Hippocampal neurons (DIV 18+) expressing mCherry were fixed in paraformaldehyde 4% in lysine phosphate buffer pH7.4 containing 2.5% of sucrose for 15-20 min. Cells were then permeabilized for 10 min in PBS containing 0.5% Triton-X 100 (Sigma). Target specific *in situ* hybridization was performed using StellarisTM probes (LGC Bioresearch) as previously described (*2*). Following fixation, cells were washed in PBS+5mM MgCl_2_, followed by dehydration in 80% ethanol overnight at −20°C. Samples were rehydrated in PBS+MgCl_2_, followed by 2x 1x sodium citrate (SSC) washes, followed by a 5 min wash in 2XSSC+30%formamide for 5 minutes. Biotin labeled probes for *Camk2a*, *Beta actin* and *Psd95* (Stellaris, Biosearch technology) were diluted into 100ul hybridization buffer and incubated with cells for 4 hours at 37°C. Following probe hybridization, samples were washed twice in 2XSSC + 30% formamide for 30 minutes each, followed by 5 1xSSC washes. After completion of *in situ* hybridization, samples were washed with phosphate buffered saline (PBS) and subsequently processed for immunofluorescence. Immunofluorescence was performed on fixed and permeabilized samples with or without *in situ* hybridization using the following protocol: samples were incubated in biotin-free blocking buffer (4% biotin free BSA in PBS) for 30 minutes and then incubated for 1½ h at room temperature or overnight at 4°C with primary antibodies in blocking buffer. After 3 washes in PBS for 5 min each, samples were incubated in blocking buffer (4% goat serum in PBS) for 1 to 2 h with secondary antibodies. The following antibodies were used: rabbit anti-biotin (Cell Signaling, 1:1000), rat anti-mCherry (Abcam, 1:1000), goat anti-rabbit Alexa 488 and goat anti-rat Alexa 568. Samples were imaged using Zeiss LSM780/880 confocal microscopes and a 63x oil objective (NA 1.4, PSF: <u>LSM780</u>- .240/.258/.729; <u>LSM880</u>- .252/.203/.563 μm x/y/z). Images spanning the entire volume of a neuron were obtained and analysed using ImageJ. To assess distance from an RNA of interest to dendritic spines, a line scan extending from the base of the nearest spine (determined by measuring the distance from an mRNA to the spines around it) through the mRNA molecule was drawn. The distance from the maximum intensity peak of the mRNA to the base of the spine was then calculated.

### FRAP translational reporters

A codon optimized superfolder GFP was custom synthesized (Eurofins) and cloned into a plasmid backbone driven by a CMV promoter. 3’untranslated (UTRs) corresponding to the most highly dendritically localized isoforms_4_ for *Camk2a*, *Beta actin* and *Psd95* were cloned upstream of a SV40 polyadenylation sequence. sfGFP reporters were transfected into neurons 12 hours prior to imaging. Cells were imaged at 63x on a LSM780 (NA 1.4, PSF: .240/.258/.729), with a temperature-regulated environmental chamber. Cells were maintained in E4 buffer_2_ supplemented with B27, Glutamax and 1x MEM amino acids (Thermofisher). Whole cell photobleaching was accomplished using a 488 argon laser (1.49mW) with an intensity of 2900kW/cm_2_ for 40-50 seconds. Cells were imaged at .067Hz for 2 minutes prior to and 10 minutes following the bleaching step. Fluorescence intensity was measured in a 50um dendritic segment from the raw image. FRAP was calculated from background-corrected fluorescence intensity by normalizing the change in fluorescence (F-F_0_) to pre-photobleaching fluorescence (F_i_). Mobile fraction and t_1/2_ values were extracted from data fitted to a one phase exponential association.

### Endogenous FRAP

Venus tagging pORANGE CRISPR/Cas9 constructs_5_ were generated from previously described GFP tagging plasmids (Addgene plasmids #131477, #131479, #131484, gifts from Harold MacGillavry). Neurons were electroporated (see above) at the day of plating and maintained until DIV17-DIV21 for FRAP experiments. FRAP and imaging was carried out using a Leica DMi8 TIRF microscope. FRAP was performed using a 488 nm laser, providing 7.78 mW/cm_2_ intensity. Cells were imaged for 2 minutes at baseline with a 488nm LED every 15 seconds prior to bleaching for a baseline measurement. Whole cell bleaching was performed with 20-30 seconds of bleaching (PSD-95) or 50-70 seconds of bleaching (CAMK2a and BETA ACTIN). Cells were the imaged in epi-fluorescence mode with the LED every 15 seconds for 60 minutes. Fluorescence intensity was measured in a 50um dendritic segment from the raw image. FRAP was calculated from background-corrected fluorescence intensity by normalizing the change in fluorescence (F-F_0_) to pre-photobleaching fluorescence (F_i_). Mobile fraction and t_1/2_ values were extracted from data fitted to a one phase exponential association.

### Puro-PLA

Detection of newly synthesized proteins by proximity ligation was performed as previously described_6_ ssing a mouse anti-puromycin (Kerafast, 1:500) antibody in combination with rabbit anti-beta actin (Abcam, 1:1000), rabbit anti-PSD-95 (cell signaling technologies, 1:1000) and rabbit anti-Camk2a (Thermo 1:1000) antibodies. PLA was performed using the duolink kit (Sigma). Rabbit PLA_plus_ and mouse PLA_minus_ probes were used for secondary antibodies along with the “Duolink Detection reagents Red” (Sigma) for ligation, amplification and label probe binding. Briefly, after a brief 5-10 minute pulse of metabolic labeling with puromycin (10uM), hippocampal cultured neurons (18+ DIV) were fixed in PBS-sucrose, permeabilized in PBS + 0.5% Triton-X 100 and blocked in PBS + 4% goat serum. Neurons were incubated overnight at 4°C in PBS + 4% goat serum containing primary antibodies. After washing, PLA probes were applied in 1:10 dilution in PBS with 4% goat serum for 1 h at 37 °C, washed several times with wash buffer A (0.01 M Tris, 0.15 M NaCl, 0.05% Tween 20) and incubated for 30 min with the ligation reaction according to the manufacturer’s recommendations in a prewarmed humidified chamber at 37°C. Amplification and label probe binding was performed after further washes with wash buffer A with the amplification reaction mixture prepared according to the manufacturer’s recommendations in a prewarmed humidified chamber at 37 °C for 100 min. Amplification was stopped by three washes in wash buffer B (0.2 M Tris, 0.1 M NaCl, pH 7.5). For better signal stability, cells were kept in wash buffer B at 4°C until imaging. Anti-Map2 immunostaining (Guinea pig anti-Map2, cell signaling, 1:5000) was performed to label dendrites. Samples were imaged using a 40x oil objective (NA 1.3, PSF: LSM780-0.217/0.260/0.566 μm; LSM880-0.238/0.253/0.636 μm x/y/z). Z-stacks (0.43 μm) spanning the entire volume of imaged neurons. Images were analyzed using ImageJ/FIJI. A 100um segment of the dendrite was assessed for the number of Puro-PLA puncta and the density of signal was calculated.

### Statistics

Statistical significance, the tests performed, and the number of cells/replicates are indicated in the figure legends. Statistical analysis was performed using GraphPad Prism.

---

**Supplemental Figure 1:**
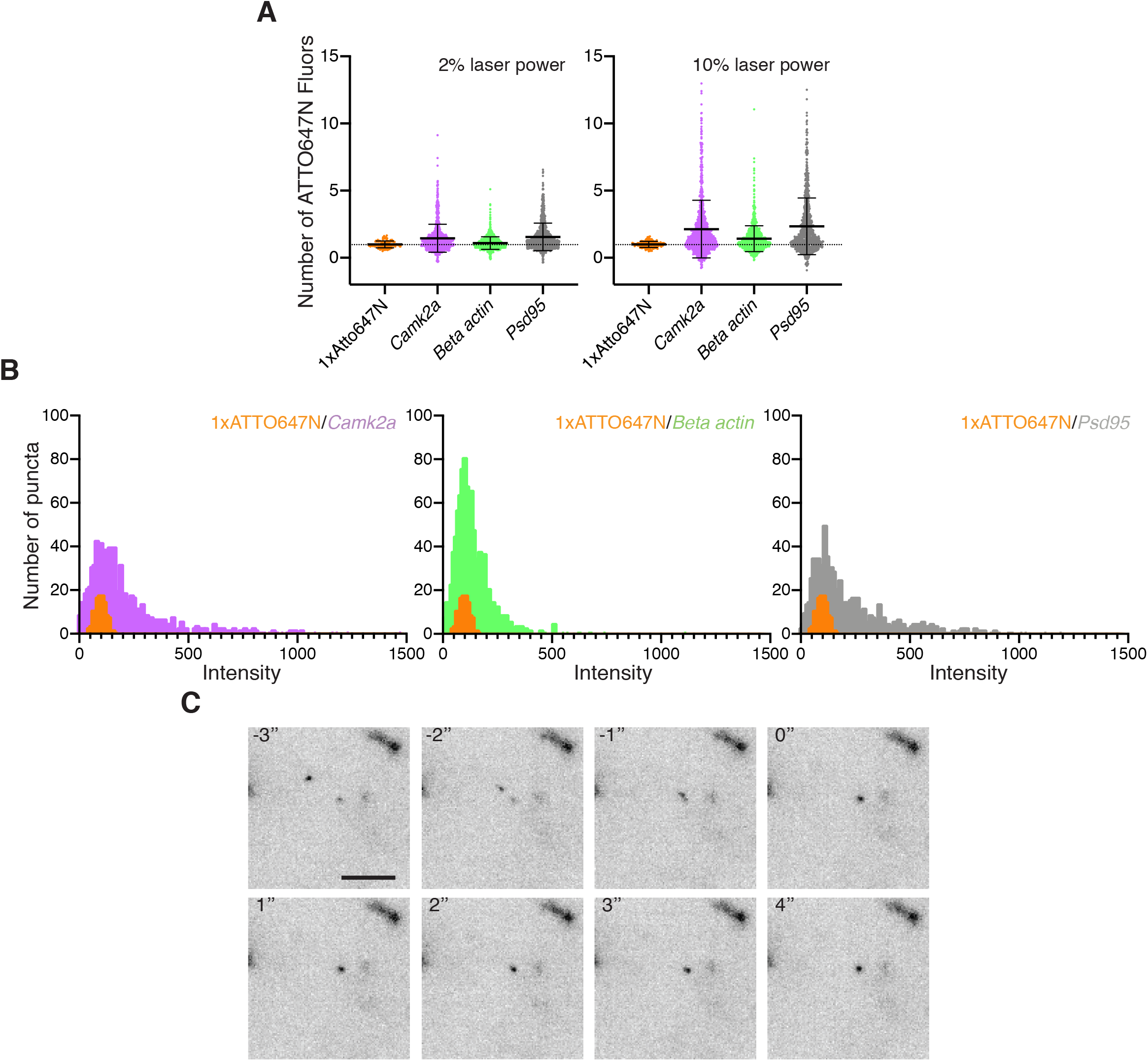
Characterization of mRNA organizational states. **A.** Quantification of mRNA organizational state within dendrites showed a heterogenous distribution pattern of copy numbers of mRNAs per granule in transcript specific manner. 1x Atto647N (3/150); *Camk2a* (19/921), *Beta actin* (21/1003), *Psd95* (18/829) (cells/number of individual mRNA granules analyzed). Comparison relative to 1xATTO647N, Dunnett’s multiple comparison test: */**** p<.05/.0001. Data shown acquired at 2 and 5% laser power, showing that the pattern of distribution holds true at various laser intensities. **B.** Plots of raw intensities of beacons and the Gatta quant standard highlighted the existence of single mRNA transport states along with higher order complex states. Data shown acquired at 10% laser intensity. **C.** Stills from a *Camk2a* 647 beacon movie (supplemental video 4) where a RNA-RNA fusion event (0”) is seen. Scale bar = 5um.

**Supplemental Figure 2:**
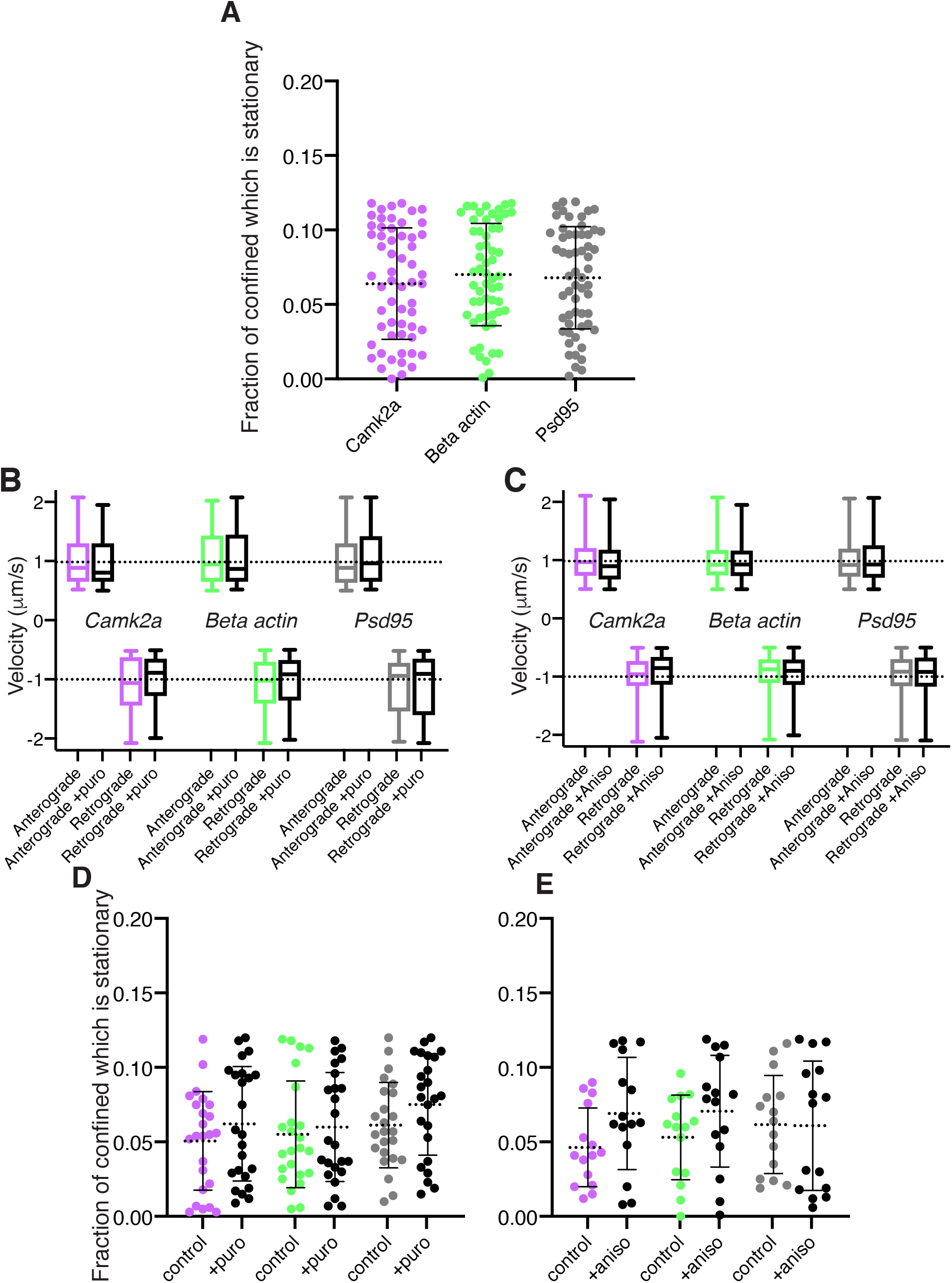
Stationary vs confined analysis for baseline and translational inhibitor experiments. **A.** Quantification of the confined population of Figure 2C which is stationary. Mean +/− SD: Camk2a 0.0640 +/− 0.0374; Beta actin 0.0701 +/− 0.0343; Psd95 0.0680 +/− 0.0344. **B.** Quantification of mRNA velocity for anterograde and retrograde control vs puro treated samples for *Camk2a (1.015 +/− 0.442 vs 0.987 +/− 0.449; −1.079 +/− .478 vs −0.991 +/− 0.420)*, *Beta actin (1.052+/− 0.489 vs 1.030 +/− 0.475; −1.130 +/− 0.449 vs −1.040 +/− 0.430)* and *Psd95 (0.988 +/− 0.414 vs 1.065 +/− 0.457; −1.111 +/− 0.484 vs −1.119 +/− .510)*. n = 29 cells per condition. **C.** Quantification of mRNA velocity for anterograde and retrograde control vs anis treated samples for *Camk2a (1.011 +/− 0.351 vs 0.9543 +/− 0.338; −0.990 +/− .335 vs −0.931 +/− 0.332)*, *Beta actin (0.993 +/− 0.334 vs 0.973 +/− 0.317; −0.942 +/− 0.308 vs −0.958 +/− 0.317)* and *Psd95 (0.990 +/− 0.343 vs 1.004 +/− 0.378; −0.973 +/− 0.340 vs −0.971 +/− .371)*. n = 15 cells per condition. **D.** Quantification of the confined population of Figure 3E which is stationary. Mean +/− SD: Camk2a 0.0507 +/− 0.0331 vs. 0.0622 +/− 0.0384; Beta actin 0.0541 +/− 0.0358 vs. 0.0599 +/− 0.0366; Psd95 0.0613 +/− 0.0287 vs. 0.0751 +-0.03402. **E.** Quantification of the confined population of Figure 3E which is stationary. Mean +/− SD: Camk2a 0.0463 +/− 0.0264 vs. 0.0691 +/− 0.0377; Beta actin 0.0530 +/− 0.0285 vs. 0.0706 +/− 0.0376; Psd95 0.0616 +/− 0.0329 vs. 0.0609 +- 0.0433.

**Supplemental Figure 3:**
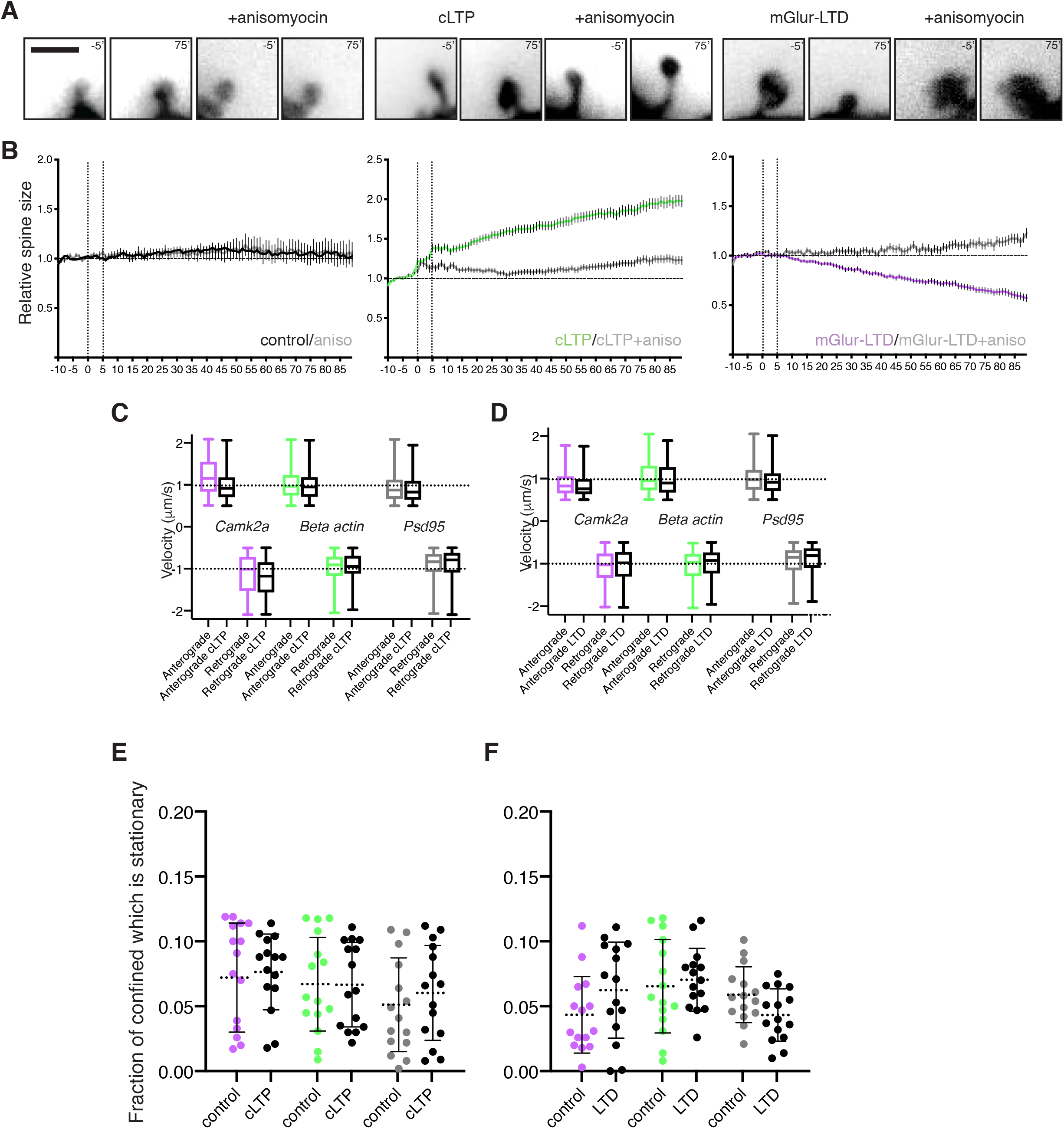
Structural plasticity changes with dendritic spines during cLTP and mGluR-LTD is protein synthesis-dependent. **A.** Example images of spines before (−5’) and after (75’) mock or plasticity stimulation in the absence or presence of anisomycin. Scale bar = 1um. **B.** Quantification of spine size over time demonstrated that cLTP and mGluR-LTD require protein synthesis for their manifestation, as spine enlargement or shrinkage is blocked by pretreatment with anisomycin. −10’-0 minutes = baseline measurement, 0-5’ = stimulation phase, 5-90’ minutes = post plasticity induction. 5-10 spines per cell were analyzed, with 20 cells in total assessed per condition. **C.** Quantification of mRNA velocity for anterograde and retrograde for *Camk2a*, *Beta actin* and *Psd95* for control vs cLTP induced samples. n = 14 cells per condition. **D.** Quantification of mRNA velocity for anterograde and retrograde for *Camk2a*, *Beta actin* and *Psd95* for control vs mGluR-LTD induced samples. n =14 cells per condition. **E.** cLTP induction did not change the overall fraction of the mRNA population which was stationary. **F.** mGluR-LTD induction did not change the overall fraction of the mRNA population which was stationary.

**Supplemental Figure 4:**
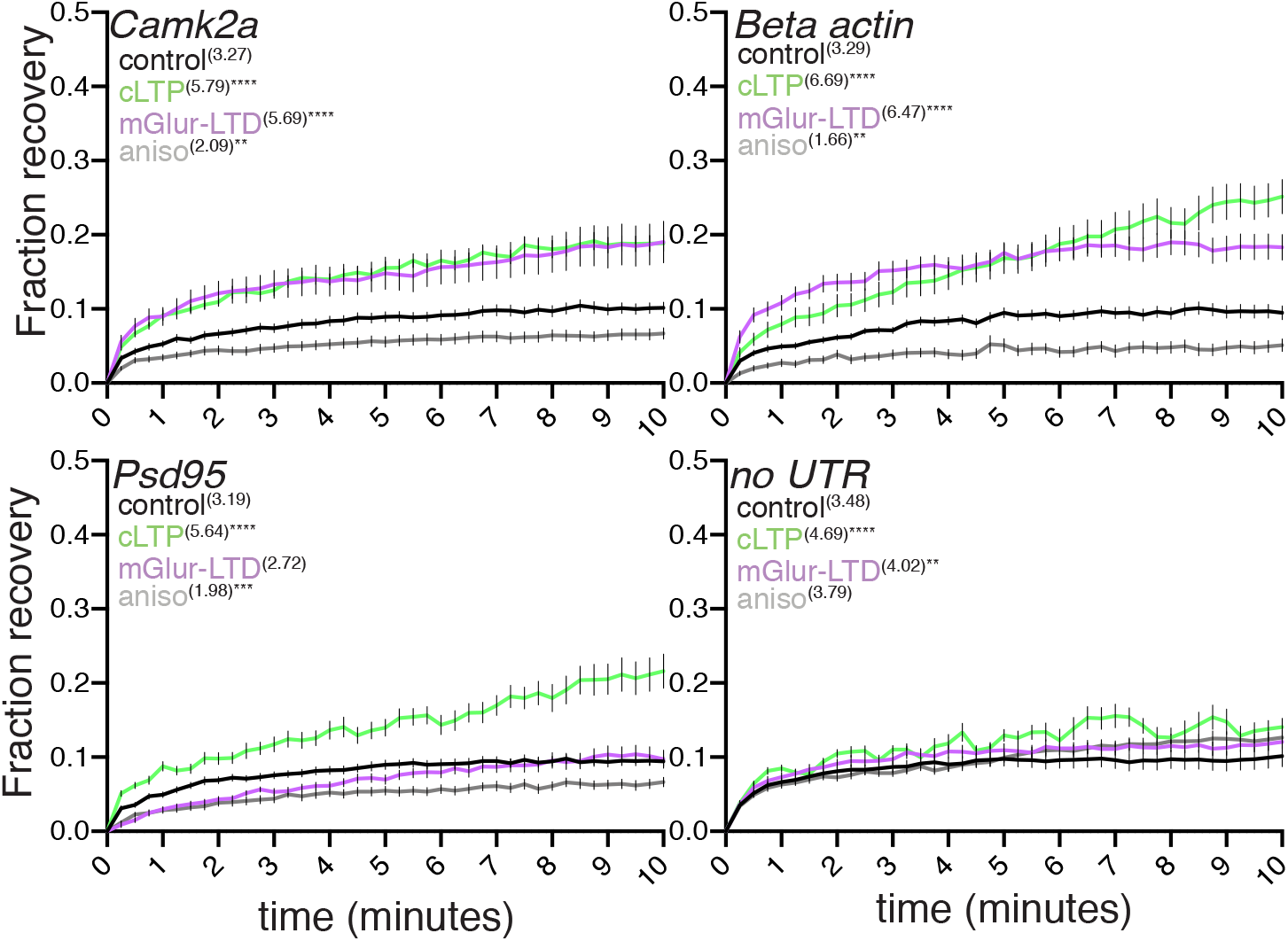
CRISPR/Cas9 labeled examples for Figure 5. Fluorescence recovery curves for all constructs under all conditions. *\*/**/***/\*\*\*\** p <.05/.01/.001/.001. Dunn’s multiple comparisons test for treated vs control condition for each construct; n = >14 cells per condition. Values in parentheses area under the curve for each condition. (area under the curve-AUC: AUC_cLTP_/AUC_control_ CAMK2a: 1.771; AUC_mGluR-LTD_/AUC_control_ CAMK2α: 1.740, AUC_cLTP_/AUC_control_ BETA ACTIN: 2.033; AUC_mGluR-LTD_/AUC_control_ BETA ACTIN: 1.967, AUC_cLTP_/AUC_control_ PSD-95: 1.768; AUC_mGluR-LTD_/AUC_control_ PSD-95: 0.852, AUC_aniso_/AUC_control_ no UTR: 1.089; CAMK2α: 0.639; BETA ACTIN: 0.504; PSD-95: 0.620)

**Supplemental Figure 5:**
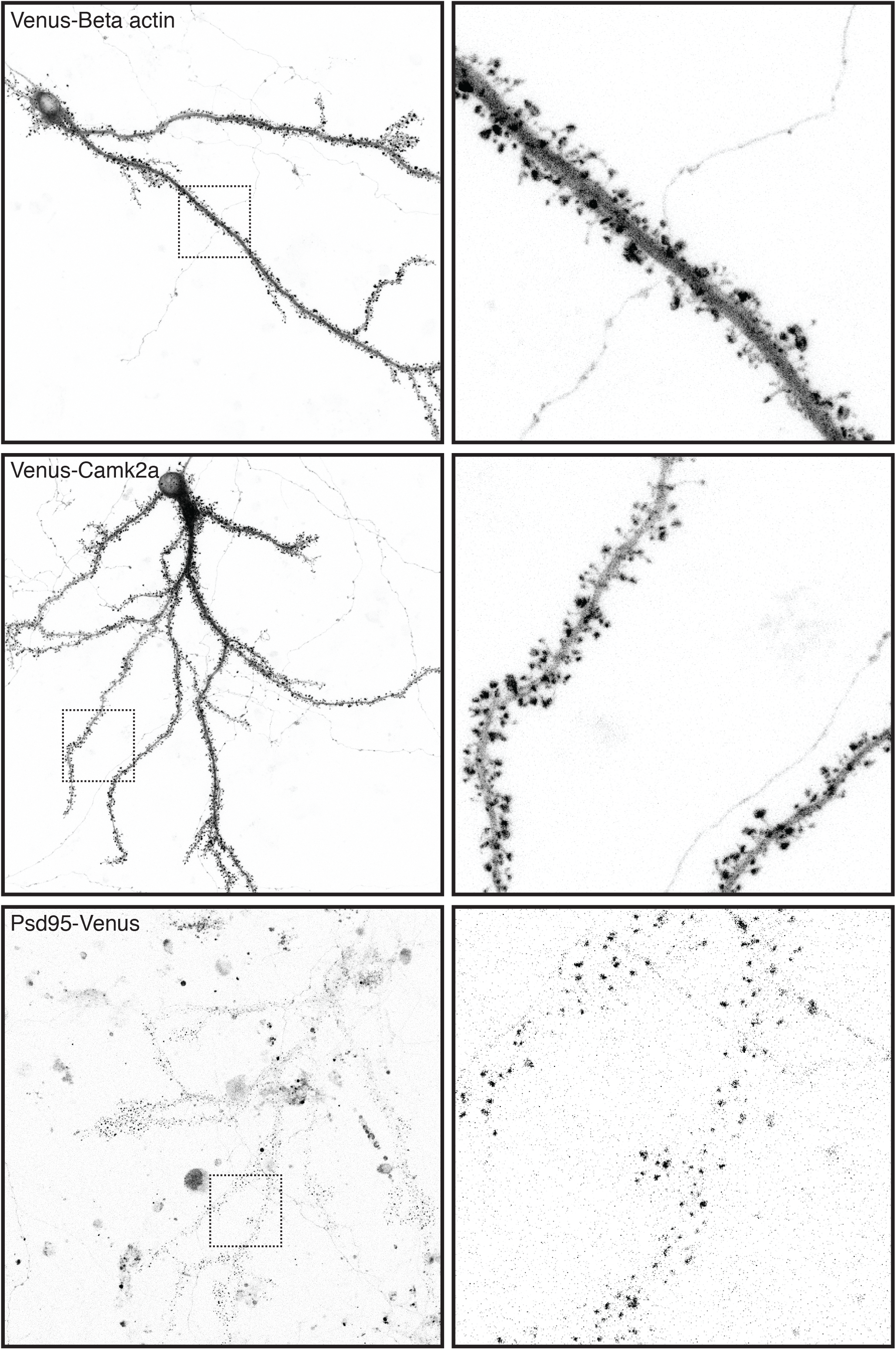
CRISPR/Cas9 labeled examples for Figure 5. **A.** Example of a Venus-BETA ACTIN labeled neuron, DIV21. Inset 40um x 40um. **B.** Example of a Venus-CAMK2a labeled neuron, DIV21. Inset 40um x 40um. **C.** Example of a PSD-95-Venus labeled neuron, DIV21. Inset 40um x 40um.

**Supplemental Figure 6:**
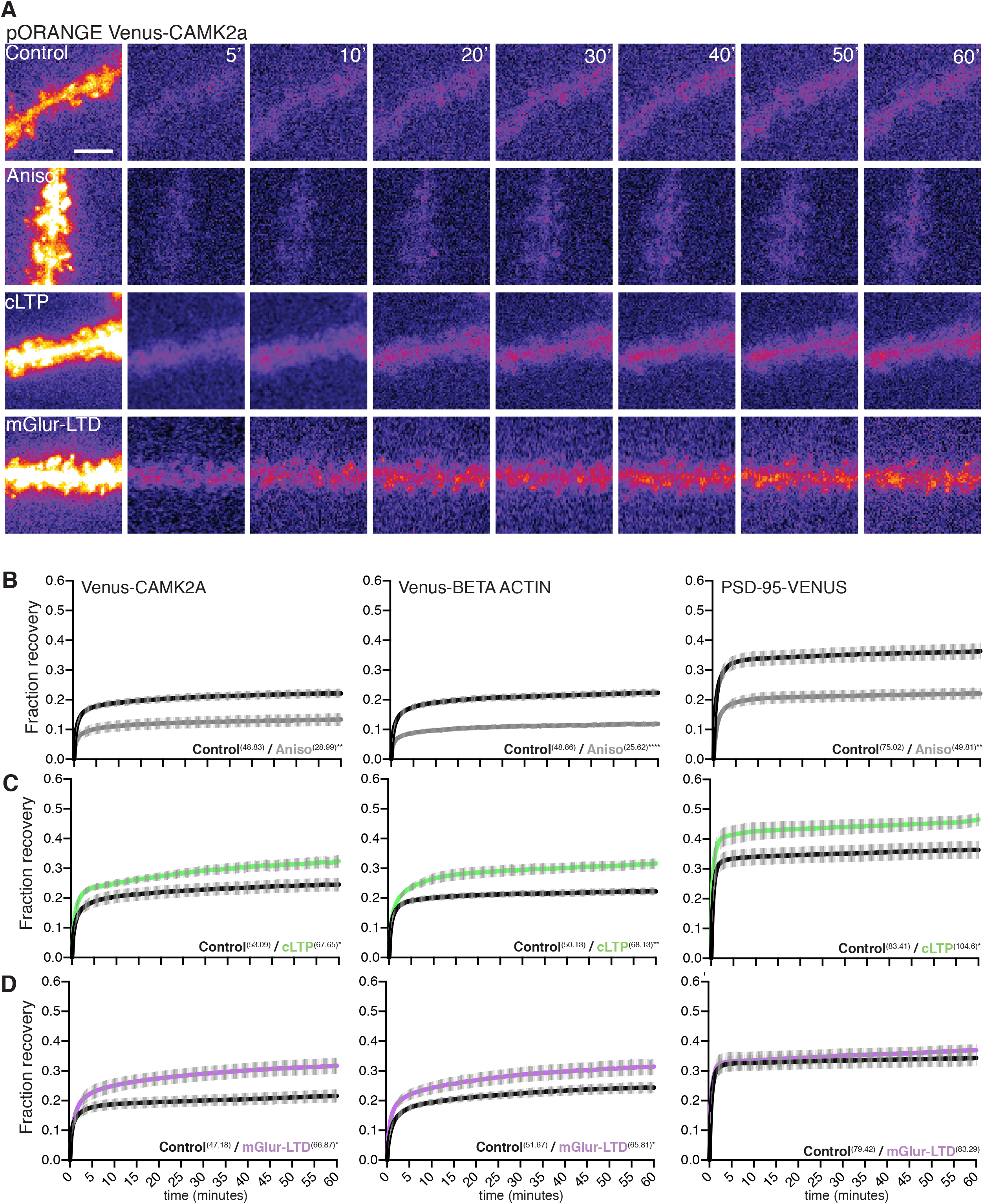
FRAP examples for Figure 5. **A.** FRAP recovery examples for Venus-CAMK2a during Control, Aniso, cLTP and mGluR-LTD. Scale bar = 5um. **B.** Fluorescence recovery curves for Venus-CAMK2a, Venus-BETA ACTIN and PSD-95-Venus with and without anisomycin treatment. *\*\*/\*\*\*\** p <.01/.0001. Dunn’s multiple comparisons test. n = 10 cells per condition. Values in parentheses are the area under the curve for each condition. (AUC_aniso_/AUC_control_ CAMK2α: 0.594; BETA ACTIN: 0.524; PSD-95: 0.664). **C.** Fluorescence recovery curves for Venus-CAMK2a, Venus-BETA ACTIN and PSD-95-Venus with and without cLTP induction. *\*/\*\** p <.05/.01. Dunn’s multiple comparisons test. n = 10 cells per condition. Values in parentheses area under the curve for each condition. (AUC_cLTP_/AUC_control_ CAMK2α: 1.274; BETA ACTIN: 1.360; PSD-95: 1.254). **D.** Fluorescence recovery curves for Venus-CAMK2a, Venus-BETA ACTIN and PSD-95-Venus with and without mGluR-LTD induction. *\*/\*\** p <.05/.01. Dunn’s multiple comparisons test. n=9-10 cells per condition. Values in parentheses are the area under the curve for each condition. (AUC_cLTP_/AUC_control_ CAMK2α: 1.417; BETA ACTIN: 1.279; PSD-95: 1.049)

**Supplemental Figure 7:**
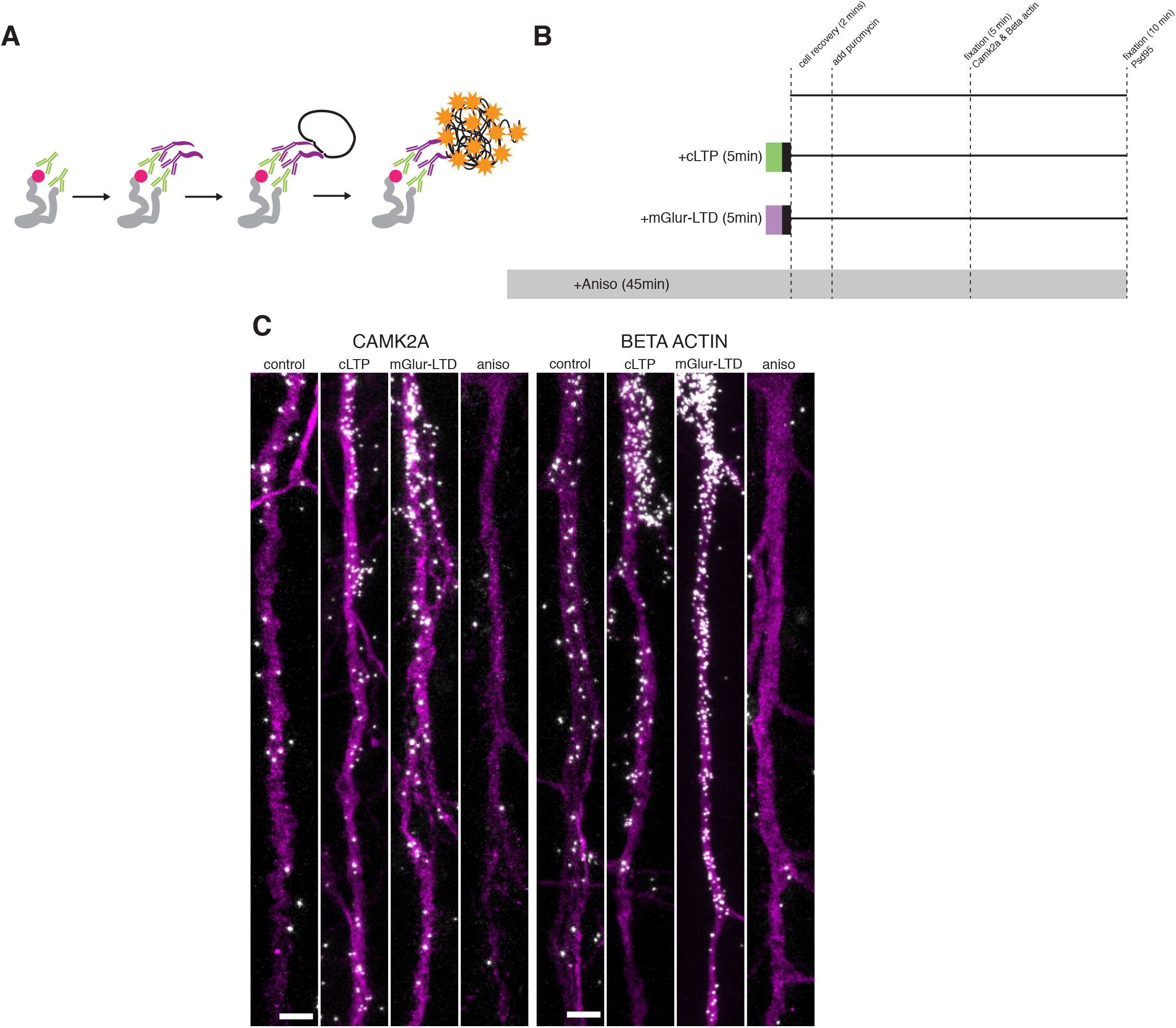
Puro-PLA examples for Figure 6. **A.** Schematic representation of the Puro-PLA method (see methods). Nascent peptides (gray) are labeled metabolically with puromycin (pink) for 5-10 minutes. A protein of interest and the puromycin tag are detected by primary antibodies (green) and then detected by PLA oligo modified secondaries (purple). With a rolling chain amplification (orange) newly synthesized proteins can be visualized *in situ*. **B.** Workflow for detecting newly synthesized CAMK2a, BETA ACTIN and PSD-95. Treatments were applied and washed out (aside from anisomycin which is maintained throughout the workflow) and cells were allowed to recover for 2 minutes prior to the addition of puromycin (10uM). CAMK2a and BETA ACTIN were fixed after 5 minutes of puromycin incubation, and PSD-95 was fixed after 10 minutes. **C.** Example images for (Figure 6F) for CAMK2a and BETA ACTIN Puro-PLA signal (gray) within dendrites (MAP2, magenta). Scale bar = 5um.

Supplemental video 1: Camk2a 647N beacon in live hippocampal neurons

Supplemental video 2: Beta actin 647N beacon in live hippocampal neurons

Supplemental video 3: Psd95 647 N beacon in live hippocampal neurons

Supplemental video 4: *Camk2a*:*Camk2a* fusion event detected in dendrites

Supplemental video 5: Beta actin example from Figure 2A&B

Supplemental Table 1: FRAP statistics for Figure 5

Supplemental Table 2: FRAP statistics for Figure 6

